# “Self-gapping” by a C-terminal domain arginine finger regulates GTP hydrolysis in bacterial zinc metallochaperones

**DOI:** 10.64898/2026.03.03.709369

**Authors:** Joseph S. Rocchio, Maximillian K. Osterberg, Emma M. McRae, Nancy Jaiswal, Katherine A. Edmonds, D. Annie Doyle, Eric P. Skaar, David P. Giedroc

**Author notes:** These authors contributed equally to this work (Joseph S. Rocchio, Maximillian K. Osterberg, Emma M. McRae). **Author Contributions:** Conceptualization, J.S.R., M.K.O., K.A.E., D.P.G.; Investigation, J.S.R., M.K.O., E.M.M., N.J., D.A.D.; Formal analysis, J.S.R., M.K.O., E.M.M., D.A.D.; Supervision, D.P.G., E.P.S.; Writing – original draft, J.S.R., M.K.O., E.M.M., D.P.G.; writing – review & editing, E.P.S., D.P.G. **Competing Interest Statement:** The authors declare no competing interests in the work described here.

## Abstract

The cellular response to transition metal scarcity is multifaceted and complex. Members of the Cluster of Orthologous Groups 0523 (COG0523) superfamily are proposed to chaperone a bound metal to activate an apoenzyme client and are thus candidate metallochaperones. COG0523 enzymes are GTPases that harbor a conserved Ras-like GTP-binding and hydrolysis domain (G-domain) and a C-terminal domain (CTD) of unknown function connected by a flexible linker. AlphaFold3 modeling posits an “open” GTPase-inactive and “closed” GTPase-active conformation where the GTP and switch 1 (G2) loop are buried at the interface of the two domains. We show here that the CTD functions as a GTP-hydrolysis activation protein (GAP) domain that stimulates GTP hydrolysis by the tethered G-domain. This “self-gapping” activity requires an invariant RxKG sequence in the β2-strand of the CTD in two distantly related bacterial COG0523s from *Acinetobacter baumannii*, ZigA and MigC. Thermodynamic and kinetic studies reveal that the Arg is analogous to the “arginine finger” motif of a Ras-cognate GAP, while the Lys residue appears to play a catalytic role in GTP hydrolysis. Cognate CTD added *in trans* to full-length RxK mutant ZigA or MigC rescues Zn(II)-activated GTPase activity whereas the non-cognate CTD shows no rescue. The linker in *Ab*ZigA appears to gate Zn(II)-stimulated GTP hydrolysis. Solution NMR studies of RxK *Ab*MigC reveal that the two domains tumble independently of one another in the absence of bound ligands, with cognate CTD added *in trans* forming a tight complex. The extent to which conformational switching characterizes eukaryotic COG0523s is discussed.

**Significance Statement:** The cellular response to nutrient transition metal limitation is evolutionarily conserved in all kingdoms, providing protection from the loss of these essential inorganic cofactors that power much of metabolism. An important part of this response is the increased cell abundance of members of the enigmatic and ubiquitous Cluster of Orthologous Groups 0523 (COG0523) superfamily. In bacterial pathogens, these enzymes are often associated with the low-zinc adaptive response to host-mediated nutritional immunity, where the host deploys transition metal chelation as an innate immune response to infections. In this work, we provide new mechanistic insights into COG0523 function with the discovery of “self-gapping” by the C-terminal domain of a two-domain G-protein architecture, placed into the context of a metallochaperone model.

## Introduction

Detection of infectious disease-causing bacteria in vertebrates triggers the immune system to release cytokines and chemokines at the site of the infection, promoting neutrophil recruitment to target areas (1). Neutrophils deploy the S100A8/S100A9 metal-chelating protein, calprotectin (CP), to sequester late *d*-block transition metal ions ranging from Mn(II) to Cu(II) and Zn(II) as part of a process known as nutritional immunity (2-6). These nutrient metals are indispensable, serving as enzyme cofactors across numerous metabolic pathways in all cells (7). *Acinetobacter baumannii*, a Gram-negative ESKAPE pathogen, displays reduced growth rates when chronically metal-starved, consistent with the observation that roughly one-third of all bacterial proteins require a metal cofactor to function (8-10). Tolerance to multi-metal restriction of Zn(II), Mn(II), Ni(II), and Fe(II) is imparted by metal-sensing transcriptional regulators (7); two that have been extensively characterized include the zinc uptake regulator, Zur, and the ferric uptake regulator, Fur (11).

*A. baumannii* exhibits a strong Fe- and Zn-starvation response when cultured in the presence of calprotectin that is characterized by increased expression of proteins that function to restore metal homeostasis and virulence (12,13). This includes the biosynthetic pathways for the catecholate siderophores acinetobactin and baumannoferrin, the Zn-uptake transporter ZnuABC, and at least four metal-sparing paralogs, including aconitase AcnB and the ribosomal protein L31B (13). ZigA, a Cluster of Orthologous Groups (COG) 0523 GTPase is Zur-regulated and accumulates to micromolar levels in cells under conditions of CP stress (13). In addition, *A. baumannii* encodes a second COG0523 protein, MigC, which while not regulated by Zur, becomes functionally important for cell wall biosynthesis under conditions of metal starvation (14). The COG0523 proteins are a poorly understood subfamily of the G3E GTPases with large knowledge gaps apparent in both enzyme structure and function (15-17). Members of the G3E GTPases include the metallochaperones UreG and HypB that are required for cofactor maturation in urease and [NiFe]-hydrogenase maturation, respectively (18-20). Consistent with a metal-delivery function, other COG0523 proteins have also been linked to cofactor maturation including the Co(II)-cobalamin maturation factor, CobW, and the Zn(II) metallochaperone for methionine aminopeptidase, Zng1 (21-24). Moreover, the Fe(II)-chaperone Nha3 in *Rhodococcus* species, which causes disease in humans and other animals, functions in the maturation of the Fe-containing nitrile hydratase, via an as yet unknown process that involves a protein-protein interaction between Nha3 and empty Fe-subunit of nitrile hydratases (25,26).

We have proposed (7) that COG0523 proteins are composed of two distinct domains: the N-terminal GTPase or G-domain which contains a family-defining and invariant CxCC (C, Cys; x, any amino acid) motif and a highly variable C-terminal domain (CTD) (14,27). The CxCC sequence donates three thiolate ligands to the high affinity, regulatory Zn(II) complex in both ZigA and MigC (14,27). Both domains are tethered by a linker that typically ranges in length from 20-40 residues and varies significantly among COG0523 proteins (17). The G-domain is a mixed α,β structure that consists of five conserved “G-loops”, G1-G5, responsible for the binding and hydrolysis of GTP into GDP and inorganic phosphate (28,29). Generally, G1-G3 stabilizes contacts with the polar phosphate groups and are intimately involved in catalysis, while G4 and G5 contribute to interactions with the nucleobase and influence nucleotide specificity. G-proteins are molecular switches where the switch 1 (G2) and switch 2 (G3) regions undergo dramatic changes in structure and/or dynamics upon GTP hydrolysis (29,30). The independently folded CTD is proposed to harbor a core αβ sandwich structure consisting of five β-strands and two α-helices, and has not been functionally characterized in any COG0523 protein (7). Notably, G3E GTPases not associated with the COG0523 subfamily, *e*.*g*., the Ni(II)-specific metallochaperones UreG and HypB, do not contain the CTD, nor do they harbor the CxCC motif, and employ a GTP-stabilized G-domain dimer to perform their functions (17). Despite these important differences, the binding of cognate metal and G-nucleotide are thermodynamically coupled in both ZigA and HypB (27,31).

A previous analysis of the COG0523 superfamily revealed an arginine and lysine residue that define an invariant “RxKG” motif, localized to the otherwise highly divergent CTDs (17) (Fig. 1). Here, we show that the RxKG motif in the two *A. baumannii* COG0523 enzymes, MigC and ZigA, is an essential functional feature. AlphaFold3 (32) modeling of ZigA and MigC carried out in the absence or presence of substrate Mg(II)•GTP or product Mg(II)•GDP and Zn(II) suggest the presence of two mutually exclusive conformations, which we term an “open” state, that mirrors the known structure of ligand-free *E. coli* YjiA (33,34) and a “closed” state not previously considered. The closed state positions an RxKG motif opposite the γ-phosphate of the bound GTP, reminiscent of the “arginine finger” motif that characterizes Ras-family GTPase activating proteins (GAPs). In the closed-state model, this motif is sandwiched between the G-domain switch I (G2) loop and the CTD. Previous G-nucleotide exchange and hydrogen-deuterium exchange-mass spectrometry studies reveal that G2 unfolds upon GTP hydrolysis (27,35).

**Fig. 1.**
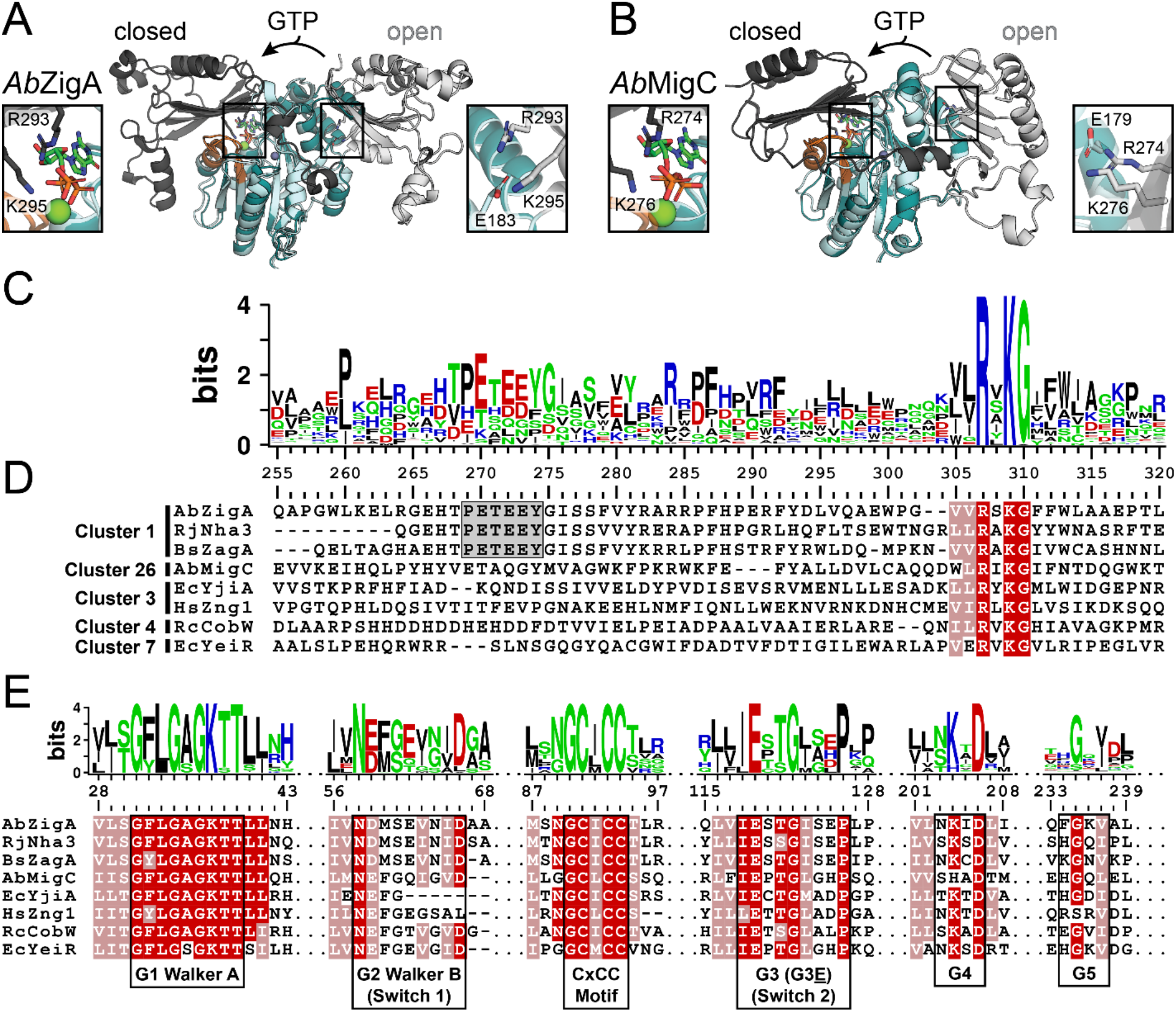
(*A-B*) AlphaFold3 model structures of (*A*) *Ab*ZigA and (*B*) *Ab*MigC modeled with no ligands (G-domains, light blue; CTD, light grey) and with Mg(II)•GTP (G-domain, dark blue; CTD dark grey; two Mg ions (green) and one Zn ion (purple) were also included in modeling). The G2-switch 1 motif is highlighted in orange in both models. (*C*) WebLogo schematic depicting sequence conservation across 363 aligned CTD sequences from functionally diverse COG0523 enzymes. (*D*) Representative multiple sequence alignment of eight previously studied COG0523 enzymes collected from the alignment. Cluster number, sequence count cluster number from the SSN analysis published previously (17), illustrated for reference. The PETEEY sequence strongly conserved in cluster 1 sequences is highlighted (*gray box*). See also *SI Appendix*, Fig. S8. (*E*) Sequence conservation of each of the five G-loops across these eight sequences, for comparison.

Here, we employ GTPase assays and isothermal titration calorimetry experiments alongside NMR solution structural studies of MigC to explore the role of the RxKG motif in nucleotide binding, metal-stimulated GTP hydrolysis and COG0523 conformational switching. These biochemical findings and biological studies with *A. baumannii* Δ*zigA* and Δ*migC* strains reveal that “self-gapping” by the RxKG motif is essential for the function of both ZigA and MigC. The implications of these findings on the mechanism of metallochaperone-dependent metalloprotein maturation, as well as to other distantly related COG0523 enzymes are discussed.

## Results

### COG0523 AlphaFold3 models predict “open” and “closed” conformational states

Structural information on COG0523 proteins is limited, with *E. coli* YjiA (*Ec*YjiA) representing the only solved structure to date (33,34). Moreover, both *Ec*YjiA structures are nucleotide-free, leaving open the question of whether these proteins can undergo conformational switching upon guanine (G) nucleotide binding. To address these limitations, we prompted AlphaFold3 (AF3) to generate structural models of ZigA and MigC, distantly related to YjiA (17), in multiple ligand-bound states (Fig. 1*A,B*) (32). Zn(II), Mg(II), and nucleotide were added to simulate the substrate-bound GTP “on” state and product-bound GDP “off” state of each enzyme (27). The GDP-bound model of ZigA is consistent with earlier AF2 structures in which the CTD is positioned opposite the nucleotide binding site (27); we designate this conformation as the “open” state. Indeed, this open-state structural model resembles the crystallographic structure of ligand-free *E. coli* YjiA (33,34). To our surprise, the GTP-bound models of MigC and ZigA both display a distinct disposition of the CTD relative to the G-domain of the open state, with the CTD placed on the opposite side of the G-domain, closer to the G-loops, and effectively burying the nucleotide at the domain interface. We refer to this modeled conformation as the “closed” state. Closer examination of both closed-state structures reveals the highly conserved RxKG motif is positioned such that these two basic side chains make direct contact with the β- and γ-phosphate groups of GTP. In contrast, models of the GDP-bound (in ZigA) or ligand-free “open” states suggest these residues form electrostatic interactions with a conserved glutamate on the opposite side of the G-domain.

### The RxKG motif is structurally reminiscent of the other GAP “arginine finger” motifs

Many eukaryotic GTPase-activating proteins (GAPs) modulate GTPase signaling using an “arginine finger” to accelerate GTP hydrolysis. This includes the well-characterized eukaryotic Ras-GAP complex, a bacterial toxin produced by *Pseudomonas aeruginosa* known as ExoS that inhibits Rac GTPase, and the heterotrimeric Gα that utilizes an arginine finger within an RxK sequence embedded in switch 1 within the G-domain (36,37). To assess the potential for similar functionality by the RxKG motif, we structurally aligned the ZigA and MigC GTP-bound closed-state models to existing GTPase-GAP complexes solved with transition-state analogues (*SI Appendix*, Fig. S1). These structures were superimposed on one another to visualize the spatial arrangement of known arginine fingers relative to RxKG motif in the ZigA and MigC AF3 models. We find that the arginine sidechain of the RxKG motif in all GTP-bound models of ZigA and MigC coincides with the arginine finger of these GAPs and supports the prediction that this conserved residue may function to stimulate GTPase activity of COG0523 family enzymes.

### The RxK motif is the only absolutely conserved sequence motif of the COG0523 CTDs

We retrieved the CTD sequences of eight (8) unique COG0523 proteins from the UniProt database previously studied. These include the bacterial enzymes, *Ab*ZigA (8,27), *Rhodococcus equi* Nha3 (25), *Bacillus subtilis* ZagA (38), *Ab*MigC (14), *Ec*YjiA (33,34), *Rhodobacter capsulatus* CobW (21), and *Ec*YeiR (21,39), and eukaryotic or plant ZNG1 proteins (22-24). These eight enzymes span a significant range of evolutionary diversity and functional understanding (17) (*SI Appendix*, Fig. S2). Using each of these eight enzymes as query in the UniProt database, we took the top ≈50 aligned sequences from each enzyme, eliminated duplicates and partial sequences, and obtained 371 unique sequences that were next globally aligned. A WebLogo plot was first generated for the first 80 residues of the CTD (Fig. 1*C*). As can be seen, the RxKG motif is invariant across this superfamily with no other superfamily-wide conservation of any sequence in the CTD. However, when one considers COG0523 enzymes most closely related to one another, it is possible to identify small regions of sequence similarity, *e*.*g*., the PETEEY sequence common to ZigA, Nha3, and ZagA, all of which reside in cluster 1 of a previous sequence similarity network (SSN) analysis (Fig. 1*D*) (17). The PETEEY motif is not found in COG0523 enzymes not associated with SSN cluster 1 (17). We also generated WebLogo plots using the conserved G-domain sequences (Fig. 1*E*). As expected, analysis of these motifs across the superfamily reveals a high degree of conservation, with the possible exception of the regulatory G2 loop (27), that contrasts sharply with the high variability observed in the CTD alignments. We conclude that like G-loops, the conservation of the RxKG motif appears an essential feature of COG0523 function.

### The C-terminal domain of MigC adopts the AF3-predicted αβ-sandwich motif

The AF3-predicted models of the CTD of ZigA and MigC reveal a mixed α,β-sandwich-like motif that features a common five-stranded antiparallel β-sheet buttressed by two α-helices, arranged in the primary structure as N-β1-α1-β2-β3-β4-β5-α2-C (*SI Appendix*, Fig. S3*A*). In the MigC model [and predicted in *Ec*YeiR (7)], AF3 predicts a sixth (N-terminal) β-strand, denoted β0, that is antiparallel to β1, thus creating a six-stranded β-sheet following the topology β0(↑)-β1(↓)-β5(↑)-β2(↓)-β3(↑)-β4(↓). The two α-helices are positioned on the same side of the β-sheet, on the outside of the closed-state structures shown in Fig. 1. ZigA differs from MigC in that there is an insertion between the β4-β5 strands into which additional helical elements may be inserted. These are positioned on the same side of the α−β-sandwich as the α1 and α2 helices, as are additional C-terminal 2º structural elements (*SI Appendix*, Fig. S3*A*). The conserved RxKG motif is found on the β2-strand, pointing in toward the G-domain.

To provide experimental support for the AF3 model of the MigC CTD, we obtained residue-specific resonance assignments for all of the backbone atoms of the core CTD (residues 232-332) (*SI Appendix*, Fig. S3*B*) and used these chemical shifts and TALOS-N (40) to predict the secondary structure of the CTD in solution (Fig. 2*A*). We find generally reasonable agreement with the AF3 model, with the exception that the β0 and β4 strands are significantly shorter experimentally; indeed, the TALOS-N predicts that the Y235-H236 dipeptide of the conserved LP**YH**Y motif (17) is not or only weakly populated in the β0-strand. The RxKG motif is indeed positioned in the β2-strand, as predicted. A semi-quantitative analysis of the ^1^H-^1^H NOESY spectra provide strong experimental support for the five-stranded antiparallel β-sheet, β1(↓)-β5(↑)-β2(↓)-β3(↑)-β4(↓), with clearly weaker support for stable β0-β1-strand packing in this construct (Fig. 2*B*).

**Fig. 2.**
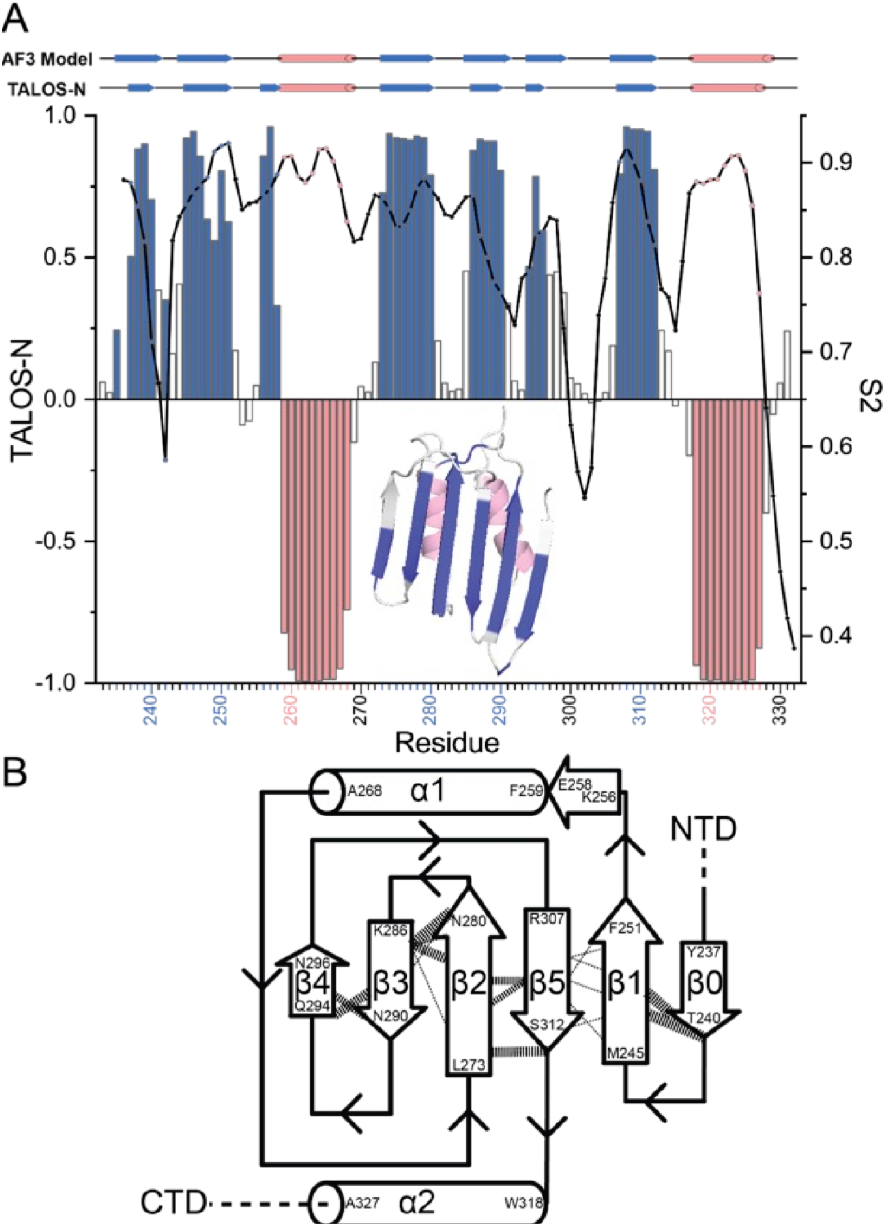
Initial characterization of the MigC core CTD in solution by NMR. (*A*) TALOS-N analysis of the core CTD (residues 233-332), schematized above the experimental data and compared to that predicted by the AF3 model. Positive TALOS-N values shaded *blue*, β-strand; negative TALOS-N values shaded *red*, α-helix; open bars, no strong prediction. Continuous black line connects the theoretical ^1^H-^15^N backbone order parameter (*S*^2^), ranging from 0 (disordered on the ps-ns timescale) to 1.0 (rigid on this timescale, tumbling with the domain itself). Note significant disorder between the β0-β1 and β4-β5 loops. *Inset*, the AF3 model shaded according to the experimental TALOS-N prediction. (*B*) Summary of key inter-β-strand NOEs mapped onto the topology predicted by TALOS-N that experimentally establishes a core five-stranded β-sheet in the MigC CTD and a weakly associated β0-strand. The width of the lines denoting the NOEs is based on the NOE quality factor calculated by CYANA (48). The PYH in the conserved ^234^**PYH**Y^237^ motif (Fig. 4*H*) is not stably folded into this β-sheet, at least not in this core CTD MigC construct.

We also obtained ^15^N-^1^H HSQC spectra for the core CTD extended on the N-terminal side to include the entire putative linker (residues 209-332), termed linker-core CTD (*SI Appendix*, S3*C*). Surprisingly, this spectrum shows sharp NH crosspeaks for many, but not all, of the assigned resonances in the core CTD, with significant line broadening specifically observed in NH groups of residues associated with the β0 and β1 strands and the structurally adjacent α2 helix (*SI Appendix*, Fig. S3*D*). In addition, two of the five Trp indole NH groups clearly visible in the core CTD spectrum (W249 in β1 and W318 in α2) are significantly broadened. Finally, the resonances associated with the entire N-terminal linker, contrary to expectations, are difficult to identify or assign in these spectra. This suggests that this region is conformational exchange-broadened as well, even in the absence of the G-domain. Clearly, extending the core CTD to include the entire linker introduces significant dynamical disorder in this region of the molecule.

### Mutations in the RxKG motif attenuates G-nucleotide binding

The AF3 modeling and sequence conservation suggests the RxKG motif contributes to COG0523 GTPase function. To investigate this, we measured apo-COG0523 binding affinities to guanine nucleotides using isothermal titration calorimetry (ITC) on mutant proteins that contain alanine substitutions at one or both positively charged residues (Fig. 3). As reported previously, WT apo-ZigA forms 1:1 complexes with GDP and GTP, exhibiting low micromolar affinities for both nucleotides with a slight preference for the product GDP (Fig. 3*A,B* *left*), findings consistent with other G3E GTPases (27,31). The R293A/K295A ZigA double mutant (RK) shows significantly weaker binding (≈60-fold) to GTP relative to the wild-type protein. The single mutant R293A ZigA (R) has a greater impact on nucleotide binding affinity than K295A (K) ZigA, ≈16-fold vs. ≈2-fold lower, respectively. The same trend is observed, albeit to a far lesser degree, with GDP. The RK double mutant binds GDP ≈16-fold weaker than WT ZigA, while R mutant ZigA shows a modest 7-fold reduction; the K mutant ZigA binds GDP with near wild-type-like affinity and energetics (*SI Appendix*, Table S1). We used a double mutant cycle analysis to measure the cooperativity between the arginine and lysine residues in G-nucleotide binding, as reflected by their pairwise coupling free energy, Δ*G*_c_ (Fig. 3*A,B*, *right*), obtained by subtracting the sum of differences in free energies of the WT ZigA and the single mutants (ΔΔG^singles^) from the difference in free energy measured for the RK double mutant (ΔΔG^double^). The Δ*G*_c_^GTP^ and Δ*G*_c_^GDP^ are –0.35 ± 0.29 and 0.07 ± 0.10 kcal mol^-1^, respectively, suggesting the mutants exhibit little, if any, energetic coupling and exert independent and additive effects on G-nucleotide binding affinity (41).

**Fig. 3.**
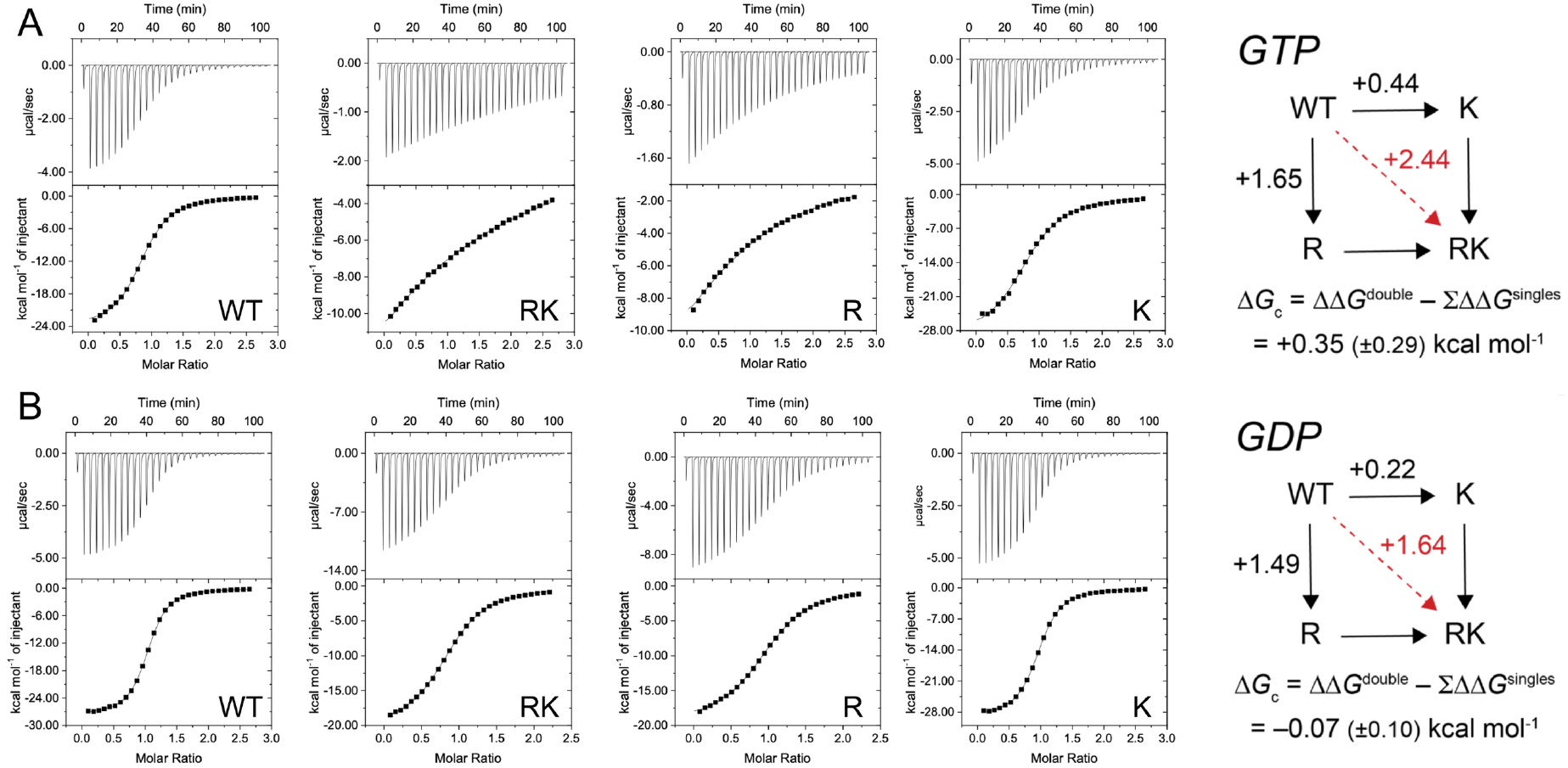
Representative thermodynamics of GTP (*A*) and GDP (*B*) binding to the apo-*Ab*ZigA as measured by isothermal titration calorimetry. Thermodynamic parameters from duplicate experiments are compiled in *SI Appendix*, Table S2.

### The RxKG motif is essential for GTPase activity

Next, we measured the GTPase activity of ZigA and MigC RxKG mutants to further characterize the extent to which these residues contribute to GTP hydrolysis. Comparisons of the conserved arginine to canonical arginine fingers (*SI Appendix*, Fig. S1) make the prediction that this residue will be important for both G-nucleotide binding (Fig. 3) *and* catalysis, while the K residue may well play an important catalytic role given its modest impact on G-nucleotide binding affinity (Fig. 3). We monitored the rate of inorganic phosphate (P_i_) release using a malachite green-based assay as a readout of enzyme turnover. Prior work establishes that ZigA and MigC activity is activated by Zn(II)-binding to the CXCC motif (8,14,35). As expected, ZigA turnover is intrinsically slow in the apo-state, and the activity increases ≈5-fold upon addition of stoichiometric Zn(II) (Fig. 4*A*, *SI Appendix*, Table S2) (35). In metal-free reactions, all three ZigA variants exhibit initial rates at or below the limit of detection; these results confirm a greatly reduced *k*_cat_ compared to wild-type apo-ZigA (Fig. 4*A*). The binding of Zn(II) to the high affinity CxCC site (*SI Appendix*, Fig. S4*A*) has no significant effect, with activities again at or below the limit of detection. These results show that the Arg and Lys residues in the RxKG motif in ZigA are required for both apo- and Zn(II)-activated GTP hydrolysis activity.

**Fig. 4.**
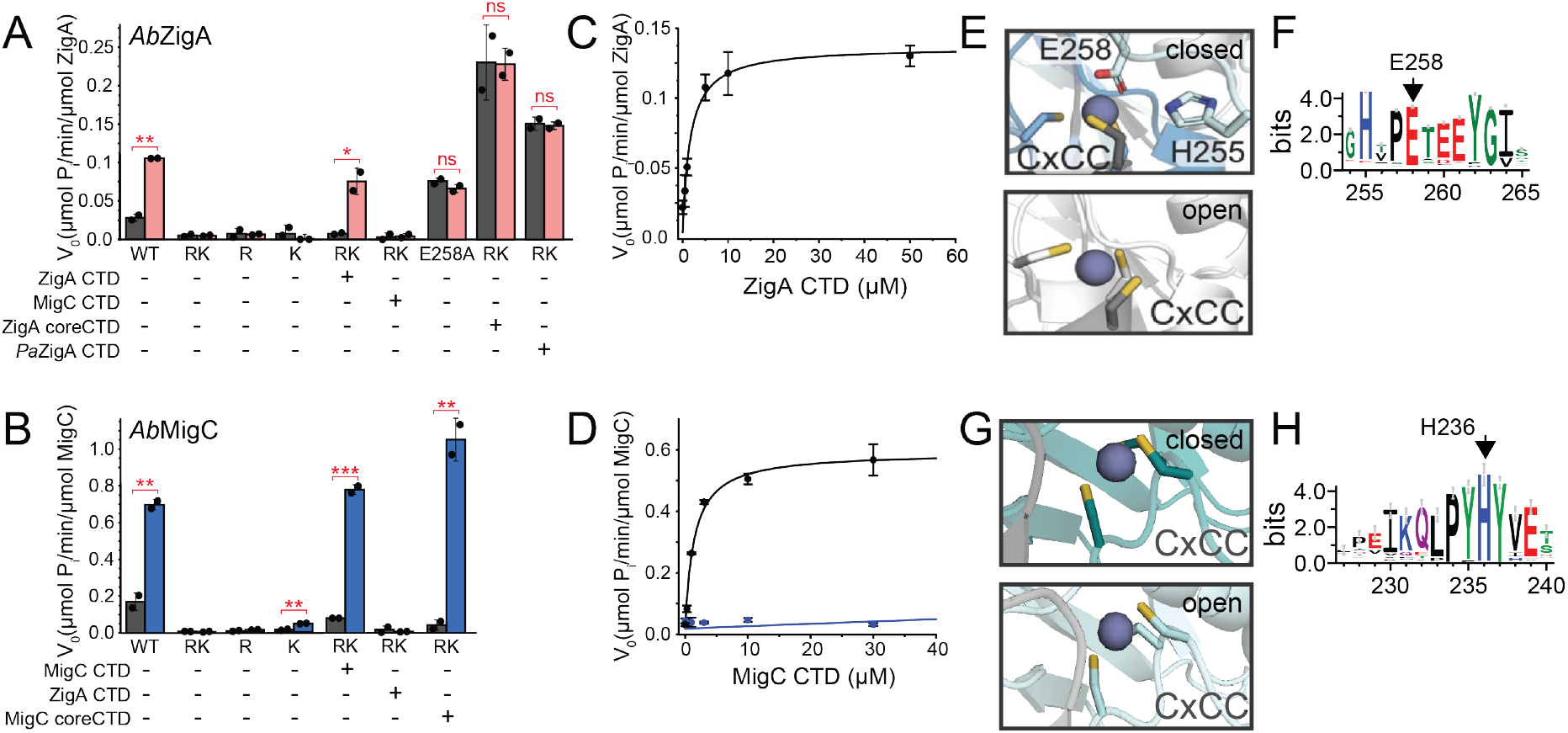
ZigA and MigC GTPase measurements. (*A*) *Ab*ZigA GTPase activity of RxK mutants (*n*=2). WT, wild-type; RK, R293A/K295A ZigA; R, R293A ZigA; K, K295A ZigA; E258A, E258A ZigA. CTD, linker-core CTD unless otherwise indicated. (*B*) *Ab*MigC GTPase activity of RxK mutants (*n*=2). WT, wild-type; RK, R274A/K276A MigC; R, R274A MigC; K, K276A MigC. CTD, linker-core CTD unless otherwise indicated. (*C*) Titration of WT linker core *Ab*ZigA CTD into RK ZigA, with the data fit to a hyperbolic function, *K*_m_=1.7 ± 0.2 µM with *V*_max_ = 0.136 ± 0.007 µmol P_i_/min/µmol RK ZigA. (*D*) Titration of WT linker core *Ab*MigC CTD into RK MigC (*black*), with the data fit to a hyperbolic function, *K*_m_=1.4 ± 0.3 µM with *V*_max_ = 0.57 ± 0.05 µmol P_i_/min/µmol RK MigC. *Blue*, linker-core ZigA CTD titration, with continuous line reflecting *K*_m_=710 ± 30 µM with *V*_max_ fixed at 0.57 µmol P_i_/min/µmol RK MigC (WT rescue value) (*n*=2). (*E*) Close-up of AF3 models around the CxCC Zn (*blue* sphere) binding region of the closed GTP-bound and open GDP-bound conformations of Zn-bound *Ab*ZigA. (*F*) Logo-plot of sequence conservation of the conserved PETEEY region in SSN cluster 1 ZigA-like COG0523 enzymes, indicating the conservation of E258 (17). (*G*) Close-up of AF3 models around the CxCC Zn (*blue* sphere) binding region of the closed GTP-bound and open conformations of Zn-bound MigC. In contrast to ZigA, the linker is not conserved and does not make close approach to the bond Zn. (*H*) Logo-plot of sequence conservation C-terminal to the linker, and just N-terminal to the β0 strand (Fig. 2) of SSN cluster 26 of MigC-like COG0523 enzymes, highlighting H236 (17). See *SI Appendix*, Table S3 for numerical values obtained for all GTPases assays.

Exactly analogous findings characterize the distantly related MigC (Fig. 4*B*, *SI Appendix*, Table S2). All three RxKG mutants, R274A, K276A and the double mutant R274A/K276A MigC are essentially inactive, regardless of the presence of Zn(II) bound to the high affinity site. These findings are collectively consistent with AF3-predicted closed-state model (Fig. 1*A*), the formation of which appears essential for GTP hydrolysis. Indeed, the increase in Zn-binding affinity mediated by GTP analog (GTPγS) binding to RK ZigA (*SI Appendix*, Table S3) is quantitatively identical to the binding of product GDP to WT ZigA (27), further support for the idea that RK ZigA cannot stably close to the closed-state and is thus inactive.

### Cognate CTD rescues the GTPase activity of the ZigA and MigC RK mutant *in trans*

We next asked if adding the wild-type CTD into these reactions could rescue RK mutant activity by supplying these catalytic residues *in trans*, mirroring the Ras-GAP mode of regulation (*SI Appendix*, Fig. S1). The C-terminal domain in ZigA is ≈20 kDa (res. 246-409; *SI Appendix*, Fig. S3*A*) and was designed to include the entire linker region between the two domains; we denote this molecule linker-core CTD. Remarkably, RK ZigA incubated with 5-fold excess cognate ZigA linker-core CTD results in significant rescue of the GTPase activity (Fig. 4*A*). Addition of linker-core to the R and K ZigA single mutants also rescues to varying degrees, but in every case, more so in the presence of Zn(II) than in the absence (*SI Appendix*, Fig. S5*A*). Finally, the non-cognate MigC linker-core CTD is inactive in this rescue activity. Varying the concentration of the ZigA linker-core CTD reveals that the *K*_m_ for rescue of GTPase activity of 1.7 µM (Fig. 4*C*).

We obtain very similar findings with the MigC linker-core CTD extended to include the linker (to T207) (Fig. 4*B*); the RK MigC is fully rescued by addition of the linker-core CTD *in trans*, with rescue again somewhat less effective for the R and K mutants (*SI Appendix*, Fig. S5*B-C*). In all cases, rescue is fully Zn-dependent. Again, the non-cognate ZigA linker-core CTD fails to rescue the GTPase activity of RK MigC at any concentration tested (Fig. 4*D*). The titration curve for the MigC linker-core CTD is hyperbolic and saturable with the *K*_*m*_ for MigC linker-core CTD of 1.4 ± 0.3 µM with a *V*_*max*_ = 0.57 ± 0.05 µmol P_i_/min/µmol (Fig. 4*D*).

These data reveal that complex formation between the cognate linker-core CTD and an intact RK mutant ZigA or MigC is specific, consistent with an extensive interface beyond addition of the catalytic residues *in trans*. These observations provide strong additional biochemical support for the closed-state model and thus “self-gapping” in both COG0523 enzymes. Marginally poorer rescue of the single R and K mutants by a wild-type CTD (*SI Appendix*, Fig. S5) might be explained by electrostatic repulsion in the assembled active site, disrupting intimate assembly of the active site around the GTP. Further, these data suggest that a wild-type linker-core CTD can readily displace an inactive intramolecular CTD, to efficiently enable “*trans*-gapping” by the cognate CTD. This is further investigated in MigC below using NMR methods.

### The linker region differentially impacts rescue by the CTD in ZigA vs. MigC

Further examination of the AF3 models (Fig. 1*A*) suggests that, in addition to the distinct orientations of the CTD relative to the G-domain, the closed conformation folds the linker over the CxCC Zn(II) binding site (Fig. 4*E*). In the specific case of ZigA, E258 from the conserved P**E**TEEY domain (Fig. 4*F*), appears to form a bidentate (*d*_Zn-O_=2.0, 2.2 Å) coordination bond to the Zn(II), to create a coordinately saturated trigonal pyramidal S_3_O_2_ chelate. We note that XAS of *Ab*ZigA loaded with GTPγS reveals a tetrahedral coordination geometry (27). In any case, this potential interaction is not conserved in MigC (Fig. 4*G*), but this is a poorly defined region (pLDDT≤50) in both AF3 models, particularly so for MigC (*SI Appendix*, Fig. S5). Remarkably, we find that an Ala substitution of the Glu residue (E258A) ZigA has greatly increased “basal” (non-Zn) GTPase activity that is no longer stimulated by Zn(II) binding to the CxCC site (Fig. 4*A*). This suggests that the linker, and perhaps an E258 coordination bond to the CxCC-bond Zn(II), plays a critical role in Zn-dependent gating of GTP hydrolysis, thus stabilizing a “locked and loaded” form of the metallochaperone (35). This finding suggests that the linker region plays a key regulatory role in allosteric coupling of Zn(II) and GTP binding observed earlier (27). This makes the prediction that the core ZigA CTD lacking the linker (and E258) may rescue RK ZigA activity in a Zn-independent fashion; this is exactly what we observe (Fig. 4*A*). Control experiments reveal that E258A ZigA binds Zn(II) with an affinity ≈6-fold more tightly as the ligand-free WT enzyme (*SI Appendix*, Table S3).

We next carried out the same series of experiments with RK MigC, determining the extent to the core cognate MigC CTD used for NMR studies stimulates MigC GTPase activity. We find that unlike the linker-core CTD MigC, GTPase activity is stimulated by core CTD to a slightly higher level than that induced by linker-core CTD and this rescue, unlike the situation with ZigA, remains strongly Zn-dependent. (Fig. 4). This reveals that the mechanism by which the GTPase activity of MigC is gated by Zn must be distinct from that of *Ab*ZigA. This makes sense since the linker does not closely approach the Zn in what is the most poorly defined region of the model, nor is this region of the linker conserved within this subclass of similar sequences (*SI Appendix*, Fig. S6) (17).

We then returned to the *Ab*ZigA system and asked if a much more closely similar linker-core CTD, derived from *P. aeruginosa* ZigA (PA5535; *SI Appendix*, Fig. S7) might rescue the *Ab*ZigA RxK mutant (Fig. 4*A*). Remarkably, we find that rescue is potent, ≈two-fold higher than cognate *Ab*ZigA linker-core CTD, but there is no Zn-dependence to the rescue. The linkers are virtually identical (31/32 residues; *SI Appendix*, Fig. S7*A*) in the two ZigAs; this suggests that optimal positioning of the tethered CTD against the cognate G-domain is required to recapitulate the full functionality of *Ab*ZigA. Indeed, the ZigA-family-specific insertion that is predicted to pack against the G-domain in the AF3 models is highly variable (*SI Appendix*, Fig. S7*B-C*, Fig. S8); this suggests that specific association of this variable region against the somewhat variable G2 loop-helix is critical for Zn-dependent gating and conformational switching.

### Conformational Changes Induce CTD binding

NMR spectroscopy was used to further characterize the WT MigC CTD and MigC mutants. First, HYDRONMR (42) modeling of the T_1_/T_2_ ratios obtained for backbone NH amides reveal that the ligand-free RK MigC behaves hydrodynamically as two largely independently folded domains (Fig. 5*A*). This suggests that the lifetime of an open state conformation (AF3 model, *inset*) must be short on the NMR time scale, with rapid relative reorientation of the two domains. This simple experiment also reveals that the individual domains of RK MigC are indeed folded, since it was straightforward to transfer the resonance assignments of the MigC core CTD (*SI Appendix*, Fig. S3*B*) to RK MigC. The behavior of wild-type MigC in the ligand-free-state is hydrodynamically larger on average than the RK mutant (Fig. 5*A*, *far left*) consistent with the idea that the wild-type enzyme samples a more compact state more often than the RK mutant. The identity of this state (open or closed) cannot be derived from these data alone, but it is likely open (RxK motif opposite E179; Fig. 1*A*), since there is no electrostatic driving force to adopt a closed state in the absence of GTP.

**Fig. 5.**
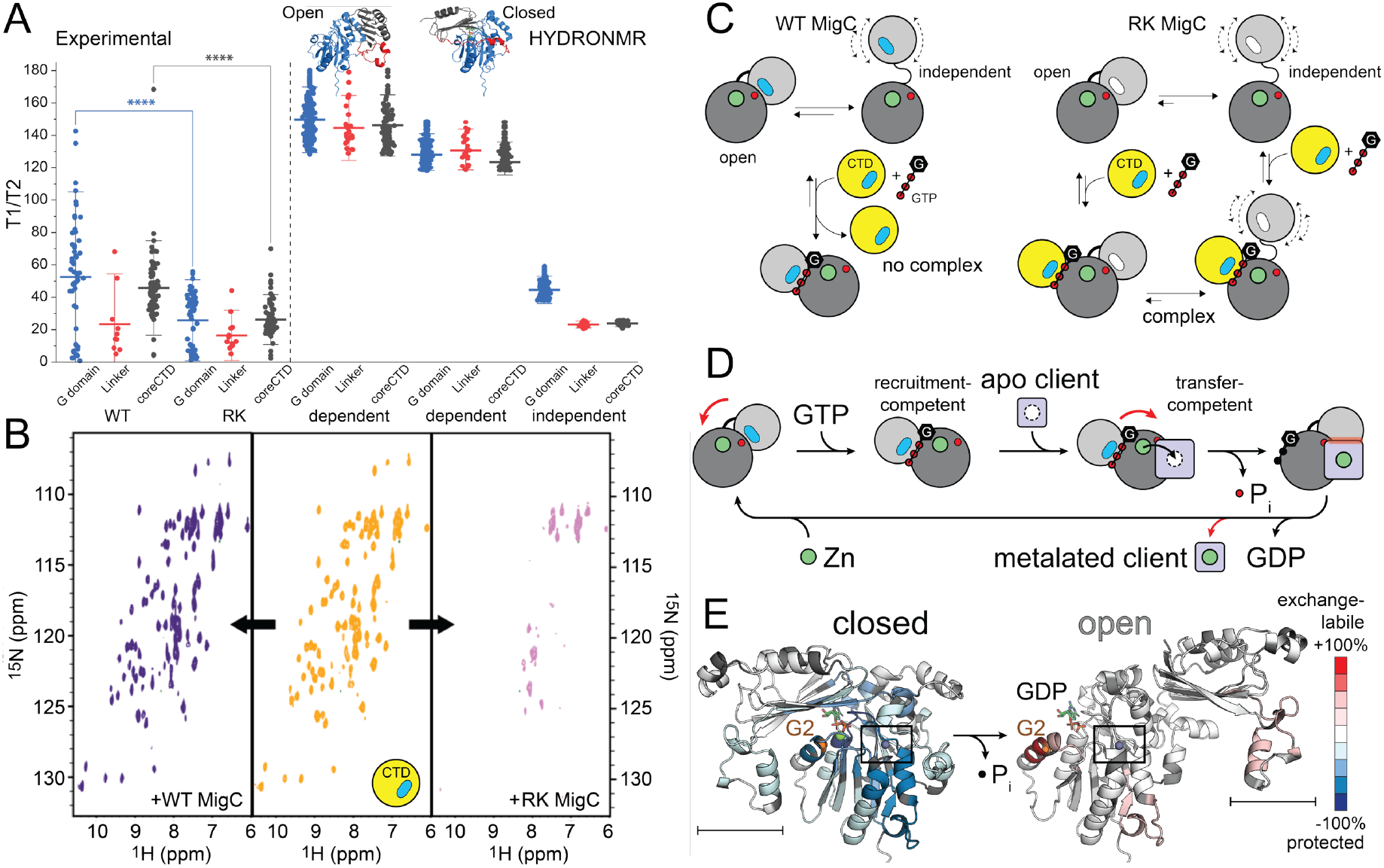
NMR characterization of CTD rescue in MigC in solution. (*A*) *Left*, experimental backbone T_1_/T_2_ data obtained for the wild-type (WT) and RK MigC in the ligand-free apo-state. *Blue*, G-domain residues; *red*, linker residues; *black*, assigned core CTD residues, indicated on the AF3 model of the open conformation, *upper right. Right*, T_1_/T_2_ values derived from HYDRONMR (42) calculated from the AF3 model of the open or closed state (dependent) vs. another model assuming independent tumbling of the G-domain and CTD (independent). ****, *p*<0.0001. The change in experimental T_1_/T_2_ for red linker residues is not significant. (*B*) ^1^H-^15^N HSQC NMR spectra of 350 μM ^15^N-labeled MigC linker-core CTD + 1 mM GDPNP (*middle*), 100 μM ^15^N-labeled MigC linker-core CTD + 1 mM GDPNP with 600 μM WT MigC added (*left*) or with 600 μM RK MigC added (*magenta*). (*C*) Cartoon rendering of these findings. MigC linker-core CTD, *yellow*; RxK motif, *blue* oval; mutated RxK motif in RK MigC, *white* oval; conserved open-state carboxylate (E179), *red* circle; Zn, *green* circle. (*D*) Metallochaperone model that takes into account the findings presented here and from (*E*) prior HDX-MS experiments with *Ab*ZigA summarized in the context of the open-closed state conformational switch proposed here (27). HDX-MS shaded according to change in protection from exchange for GTPγS–apo states (*left*), GDP–GTPγS states (*right*) according to the scale bar, *right*. The ZigA-specific insertion is encompassed by the horizontal bar (see also *SI Appendix*, Fig. S7-S8), which enfolds the G2-switch 2 motif only in the closed-state AF3 model. The Zn coordination site (Zn, *purple* sphere) is boxed in each model, for reference.

We next asked if these conformational dynamics could impact the binding of the linker-core CTD to the intact enzyme. We first acquired a reference ^1^H-^15^N HSQC spectrum of ^15^N WT MigC linker-core CTD (Fig. 5*B*, *middle*) and then added intact WT MigC to a roughly two-fold molar excess (Fig. 5*B*, *left*). As can be seen, no spectral perturbations are observed. In contrast, when RK MigC is added, we observe nearly complete loss of all backbone signals indicative of complex formation (Fig. 5*B*, *right*). This complex may well be in intermediate chemical exchange with the free CTD, consistent with the low µM *K*_d_, which is anticipated to give rise to significant chemical exchange broadening if one assumes a reasonable on-rate (43). These data, taken collectively are consistent with the model shown (Fig. 5*C*), which suggests that MigC can populate a number of distinct conformation states that are dependent on the integrity of the *cis* RxKG motif in the CTD. These features may well be important mechanistically during the projected metal transfer process (Fig. 5*D*) and are fully consistent with previously published HDX-MS experiments of *Ab*ZigA carried out prior to consideration of a closed, “self-gapped” conformation of the metallochaperone (Fig. 5*E*) (see Discussion).

### Mutations of the RxK motif inhibit ZigA and MigC function *in vivo*

Previous work reveals that *A. baumannii* strains that lack the *zigA* or *migC* genes have a growth defect when grown in transition metal restricted conditions, *e*.*g*., in the presence of the physiological chelator calprotectin or the broad-spectrum cell-permeable transition metal chelator, TPEN (*N,N,N′,N′-* tetrakis(2-pyridinylmethyl)-1,2-ethanediamine) (8,12,14). Here we extend these data to assess the extent to which complemented RxK mutant alleles can rescue the TPEN-induced growth phenotype of the corresponding deletion strain compared to a wild-type allele as a measure of functionality of ZigA (Fig. 6*A*) or MigC (Fig. 6*B*) in *A. baumannii* cells. All strains grow like WT in the absence of TPEN and exhibit growth inhibition when exposed to a TPEN in a concentration-dependent manner (*SI Appendix*, Fig. S9*A-C*, with the results shown at the highest TPEN concentration shown in both cases (Fig. 6). The TPEN-induced growth phenotype of *A. baumannii* Δ*zigA* is rather modest and is largely complemented by expression of the WT allele. However, the R, K and RK *zigA* complemented mutants appear to behave largely like the deletion strain in this assay, suggesting that substitution of each residue within the RxK motif results in poor functionality within the cell (Fig. 6*A*).

**Fig. 6.**
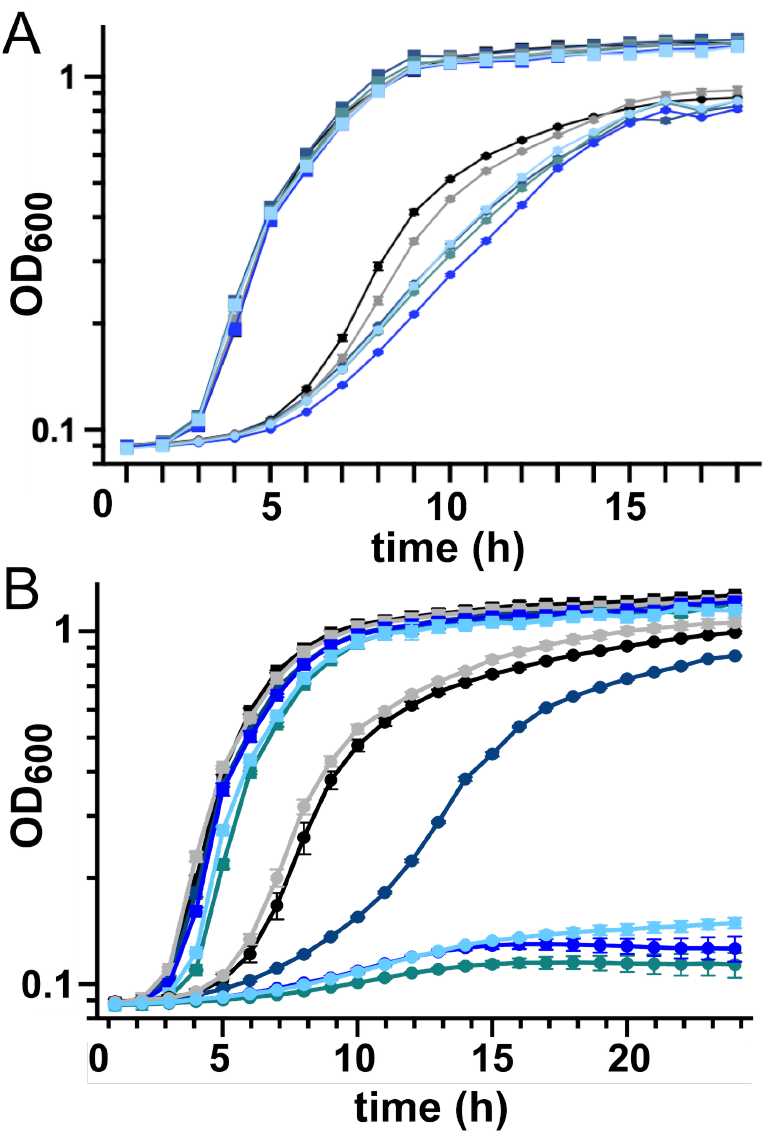
Growth curves for various *A. baumannii* strains with and without added TPEN. (*A*) Δ*zigA* strains chromosomally complemented with the indicated *zigA* allele compared to the parent wild-type (WT) strain grown in the presence (*circles*) of 80 µM TPEN vs. the same amount of ethanol used to dissolve the TPEN (–TPEN; *squares*). WT (black), Δ*zigA* (dark blue), Δ*zigA*:*zigA* (gray), Δ*zigA*:*zigA*_R293A_ (blue-gray), Δ*zigA*:*zigA*_K295A_ (blue), Δ*zigA*:*zigA*_R293AK295A_ (cyan). (*B*) Δ*migC* strains harboring a pWH1266 complementation vector with the indicated *migC* allele compared to the parent wild-type (WT) strain grown in the presence 40 µM TPEN (*circles*) vs. no addition of TPEN (–TPEN; *squares*). WT (black), Δ*migC* (dark blue), Δ*migC:migC* (gray), Δ*migC*:*migC*_R274A_ (blue-gray), Δ*zigA*:*zigA*_K276A_ (blue), Δ*zigA*:*zigA*_R274AK276A_ (cyan). See *SI Appendix*, Fig. S9 for other companion growth curves with these strains.

The analogous experiment with *A. baumannii* Δ*migC* and complemented strains is even more striking (Fig. 6*B*). Again, there is little to no growth phenotype in the absence of TPEN in any strain, and the significant growth phenotype of the Δ*migC* strain is fully complemented by expression of the WT allele. Expression of R, K and RK mutants fails to complement but gives rise to strong inhibition of growth (see also *SI Appendix*, Fig. S9*E*). This finding suggests a dominant-negative effect, in which a *migC* allele unable to catalyze GTP hydrolysis may further inhibit the biological process in which MigC participates, perhaps by binding to and inhibiting MigC client enzyme(s) activity. These biological studies reveal that the RxK motif in both ZigA and MigC are required for full functionality in cells.

## Discussion

The findings reported here provide new insights into functional characterization of the two COG0523-family metallochaperones in *A. baumannii*, ZigA and MigC. Given the limited structural information available, AlphaFold3 models allowed us to propose G-nucleotide-selective open and closed conformational states that may well broadly characterize this family of enzymes. The states differ by a striking migration of the C-terminal domain from one side of the G-domain to the other, with a very clearly defined “pivot” point in both systems (*SI Appendix*, Fig. S6). An important feature of the closed state structures is the spatial arrangement of the invariant RxK motif derived from the CTD around the bound GTP, coinciding with previously characterized “arginine fingers” in eukaryotic Ras and Ras-family enzymes that stimulated GTP hydrolysis. The R and the K residue play distinct roles in GTP binding and catalysis. An Ala substitution of the R residue in both ZigA and MigC strongly impacts GTP binding, while the impact of Ala substitution of the K residue has a comparatively smaller effect; however, both the R and K residues are essential for catalysis.

We were surprised to see that an Ala substitution of R and/or K residue in ZigA negatively impacts product GDP binding as well. This suggests the possibility that specific G-nucleotide binding (GTP vs GDP) shifts a pre-existing open-to-closed conformational equilibrium toward one or the other state, rather than inducing a conformational switch from purely one state to another. Indeed, the ligand-free RK MigC appears to behave hydrodynamically as though the two domains are tumbling independently of one another; this is also true of the wild-type enzyme, albeit not to same extent. In addition, the cognate CTD added *in trans* is capable of binding to the RK MigC, perhaps pushing the *cis*-domain aside, but not in the wild-type enzyme. These data suggest the RxK-mediated electrostatic interactions with the G-domain play a central role in driving conformational switching, particularly in the GTP-hydrolysis-competent closed state.

The finding that the RK mutant GTPase activity of both ZigA and MigC is rescued in a Zn-dependent fashion by the cognate linker-core CTD *in trans* has broad implications regarding interdomain dynamics and allostery in COG0523 superfamily enzymes. These interdomain interactions are supported by the single-digit µM *K*_m_ estimated for linker-core cognate CTD activation. It is interesting to note that the activity of the *trans*-reconstituted enzyme in the absence of Zn is far less than the intact enzyme; nonetheless, Zn-dependent GTP hydrolysis of the rescued RK mutant is comparable to, or greater than the intact enzymes. This suggests that linker itself is not passive but rather plays a regulatory role in CTD rescue, since the core CTD-stimulated GTP hydrolysis activity is distinct from that of the linker-core CTD. We anticipate that these *trans* features characterize the “self-gapped” intramolecular native-state conformation as well.

Close examination of the CTD rescue activity by linker-core vs. core CTDs in RK ZigA vs. RK MigC reveals clear differences. In the case of ZigA, rescue by the core CTD becomes independent of Zn (and higher in activity), even while retaining the high affinity CxCC Zn site in RK ZigA (*SI Appendix*, Fig. S4 and Table S3). This may be due solely to the loss of E258 in a conserved region of the linker (*SI Appendix*, Fig. S7), since in this mutant, Zn-dependent gating of the GTP hydrolysis is also lost. Interestingly, this mutant retains the high affinity CxCC Zn-binding site, but Zn binding here is no longer strongly coupled to G-nucleotide binding (27). Perhaps E258 provides a fourth non-thiolate ligand to Zn coordination complex in the “on” state. However, rescue by *Pa*ZigA reveals that E258 alone is not sufficient to enable Zn-dependent regulation of self-gapping in ZigA, with regions beyond the linker clearly playing a critical role. It is tempting to speculate that full regulatory activity this involves an intimate association of the ZigA-specific insertion (*SI Appendix*, Fig. S7*C*) with the G-domain that effectively surrounds the G2-switch 1 determinants that unravels upon GTP hydrolysis, thus providing a driving force to pivot to the open, GDP-bound “off” state (Fig. 5*D-E*) (27). In any case, the linker helps to ensure high GTP hydrolysis activity only when the Zn cargo is bound by the metallochaperone.

In contrast, the linker is not required to gate GTP hydrolysis in MigC by Zn, suggesting that the mechanism of Zn-dependent activation of “self-gapping” is distinct in these distantly related metallochaperones. In the case of MigC, the conserved region lies only in the region just upstream of the short β0 strand (PY**H**Y) (Fig. 4*H*), with this sequence not part of this β-strand in solution. H236 in this sequence is an excellent candidate for playing a regulatory role analogous to E258 in ZigA. Indeed Zn-XAS studies strongly suggest the presence of a His residue as the non-thiolate Zn ligand in MigC (14). This has not yet been tested. In any case, these findings seem to suggest the considerable sequence diversity observed in COG0523 C-terminal domains would represent an effective evolutionary strategy to avoid crosstalk among multiple members of this superfamily of putative metalloenzyme activators operating in the same cell (44).

The connection identified here between the RxK arginine in our metallochaperone models and GAP arginine fingers in other GTPases can now be placed into context of the canonical GTPase signaling cycle between an active “one” and inactive “off” state. The arginine finger observed in COG0523 proteins suggests a “self-gapping” signaling cycle like that of heterotrimeric G-proteins, where the arginine finger from switch 1 is positioned into the active site to stabilize the transition state and accelerate GTP hydrolysis. GAPs position the arginine finger to neutralize the negative charge generated on the γ-phosphate and stabilize the transition state during hydrolysis, resulting in rates many times greater than the intrinsic activity (45). Our AF3 models suggest that the RxK motif arginine donated by the CTD induces a similar electrostatic stabilization of the transition state in both ZigA and MigC, which can also be provided *in trans*. The function of the conserved RxK lysine residue, on the other hand, is less clear. A conserved glutamine derived from the G3/switch 2 loop in at least a subset of GAPs may well be responsible for activating a water molecule that performs the nucleophilic attack onto the γ-phosphate (46). Other GAPs, however, do not conserve this glutamine, and therefore must employ other mechanisms to lower the energy of the transition state and activate a water molecule for catalysis (46). Inspection of the AF3 model suggests that the only residue near the γ−phosphate that might perform such a role is the lysine in the RxK motif; however, its p*K*_a_ would have to be more acidic than a typical Lys side chain to function as a catalytic base in activating a water molecule. The close physical placement of the Arg and Lys side chains may well enforce lower p*K*_a_ of K295 to minimize electrostatic repulsion of the two positively charge side chains in the transition state. Such a role would be consistent with the finding the K295A mutant shows only a minimal impact on guanine nucleotide binding affinity but is catalytically inactive.

In light of these findings and those published previously (27), we present a revised model for metallochaperone function (Fig. 5*D-E*) where conformational switching between GTP and Zn-bound closed state and an open GDP-bound state are readily integrated. Here, the closed state buries GTP at the interdomain interface, consistent with the kinetic trapping of GTP (but not GDP) observed earlier, for ZigA (35). We refer to this state as a client enzyme “recruitment-competent” state that is ready to engage client. We note that the G2 loop-helix forms part of the interface in the closed state; GTP hydrolysis unravels this helix, at least in *Ab*ZigA (27), which may contribute to domain migration while also providing a favorable thermodynamic gradient for metal transfer (27). Domain migration in the GDP-bound state then alters the interface with metalated client, leading to disassembly of the complex.

The extent to which these proposed structural and mechanistic features are operative for either ZigA and MigC is unknown and must await the identification of a *bona fide* client enzyme(s) for each metallochaperone. The use of an *A. baumannii* strain harboring the RK mutant allele may well facilitate this (Fig. 6). The extent to which these features characterize other distantly related COG0523 enzymes, including eukaryotic ZNG1 and its metalloenzyme client, methionine aminopeptidase-1, is also unknown (23,24). AF3 modeling of vertebrate ZNG1 in the GTP vs. GDP bound states suggests a very similar conformational switch (not shown), although how this impacts ZNG1 metal delivery to METAP1 and other clients remains unknown. An exciting future avenue of investigation will be to work toward structural characterization of a trapped client protein-metallochaperone complex through use of a ground state or transition state analog, *e*.*g*., GDP•BeF_3_ or GDP•AlF_3_, respectively, that mimics the structure of the post-metal transfer complex, but prior to P_i_ release and potentially, conformational switching to the open state (Fig. 5*D*) (47). These and other studies are underway in our laboratories.

## Materials and Methods

### AlphaFold3 modeling

AlphaFold3 models were generated on the online Alphafold Server (https://alphafoldserver.com). Sequences for *Ab*ZigA (A0A7U4X7Y7), *Ab*MigC (A0A0D5YLB1), and *Pa*ZigA (locus tag PA5535; Q9HT39) were simulated with one equivalent of GTP, Mg(II), and Zn(II) to observe the “closed” state. For the “open” state, *Ab*ZigA and *Pa*ZigA were run with GDP, Mg(II), and Zn(II), while *Ab*MigC was run with Zn(II) only. Five models were generated for each and the highest ranked model, as determined by PAE and pLDDT scores, was considered the representative structure. The seed for each prediction was as follows: *Ab*ZigA open (940640380), closed (1094485029); *Ab*MigC open (2122444129), closed (1124575016). These models are available at ModelArchive (https://modelarchive.org) with the accession codes: ma-iwqsx, *Ab*MigC bound to Mg(II)•GTP and Zn(II); ma-l1a07, *Ab*MigC bound to Zn(II); ma-h76qi, *Ab*MigC bound to Mg(II)•GDP and Zn(II); ma-tca8p, *Ab*ZigA bound to Mg(II)•GTP and Zn(II); ma-bzf37, *Ab*ZigA bound to Mg(II)•GDP and Zn(II); ma-d36dv, *Ab*ZigA bound to Zn(II).

### GTPase Assays Monitoring ZigA and MigC Activity

Reactions measuring MigC and ZigA GTPase activity were performed in 25 mM HEPES, 150 mM KCl, 2 mM MgCl_2_, 2 mM TCEP, 500 μM GTP, pH 7.4. Zn containing MigC or ZigA was pre-loaded prior to assays with 1 mol equivalent of ZnSO_4_. Rescue reactions containing MigC or ZigA linker-core and core CTDs were run at a single concentration of 30 μM. Reactions (100 μL) were initiated with the addition of 5 μM protein and incubated at 37 °C for 90 min. Ten μL aliquots were removed at specific times (typically 2 min, 15 min, 30 min, 45 min, 60 min, and 90 min; shorter time periods for faster rates) and quenched with 100 mM EDTA. Inorganic phosphate was detected with the addition of 80 μL of 1 mM malachite green oxalate (Acros Organics), 10 mM ammonium molybdate (Sigma-Aldrich) in 1 M HCl. This sample mixture was allowed to incubate in the dark at ambient temperature for 10 min and then quenched with the addition of 10 μL 35% citric acid. Mixtures were incubated for 10 min and inorganic phosphate concentration was calculated based on the absorbance at 660 nm relative to a standard curve (5–500 μM inorganic phosphate, Fluka Analytical). The production of inorganic phosphate over this time period was linear and used to calculate the initial rate for each reaction (V_o_) in units of μmol P_i_ min^−1^ μmol enzyme^−1^. A rate of ≤0.0002 μmol P_i_ min^−1^ μmol enzyme^−1^ was assigned a rate of 0, as the lower limit over this assay. The measured rates represent an average of two independently performed reactions (*SI Appendix*, Table S2).

Other methods are described in the ***Supporting Information***.

## Acknowledgments

This work was supported by NIH grants R35 GM118157 (to D.P.G.), R01 AI101171 (to E.P.S. and D.P.G.) and R01 AI178929 (to E.P.S and D.P.G.). M.K.O. and E.M.M. were supported by graduate fellowships provided by the training program in Quantitative and Chemical Biology (T32 GM131994). D.A.D. was supported by NHLBI training grant T32 HL094296.

