## Supplemental Material for "“Self-gapping” by a C-terminal domain arginine finger regulates GTP hydrolysis in bacterial zinc metallochaperones"

This file contains Supplementary Methods, Supplementary Tables S1-S4, and Supplementary Figures S1-S9 and Supplementary References.

### Supplementary Methods

**Protein Expression and Purification.** The gene *a1s_0934* encoding MigC was amplified from *A. baumannii* ATCC 17978 genomic DNA using *MigC_pHIS_F* and *MigC_pHIS_R* (primers listed in *SI Appendix*, Table S4) and cloned into pHIS-Parallel1 expression vector at the NdeI restriction site (1) as described earlier (2). MigC expression and purification was carried out as described, confirmed to be metal-free by ICP-MS, with protein concentrations measured by UV-Visible spectroscopy using $\varepsilon_{280}$ of 62,910 $M^{-1}$ $cm^{-1}$ (2). Mutagenesis was performed on pHIS-Parallel1 expression vector containing *migC* using the primers listed (*SI Appendix*, Table S4). All MigC point mutants were purified as described for the wild-type enzyme with only minor modifications. Wild-type *Ab*ZigA was expressed and purified as described previously (3). Mutagenesis was performed using the primers listed (*SI Appendix*, Table S4) and pHIS-Parallel1 expression vector containing *zigA* as template, and each mutant was purified and characterized as described with the wild-type enzyme. All *Ab*ZigA mutants were confirmed to be metal-free by ICP-MS.

Expression and purification of linker-core and core *Ab*ZigA and *Ab*MigC C-terminal terminal domains was carried out largely as described for *Ab*ZigA (3). The gene encoding *P. aeruginosa* PAO1 PA5535 (hereafter *Pa*ZigA) was cloned using PaZigA_pHis_F and PaZigA_pHis_R primers (*SI Appendix*, Table S4) into the pHIS-Parallel1 expression vector, expressed and purified using a protocol developed for *Ab*ZigA with minor modifications. *Pa*ZigA linker-core CTD (residues W293 to A400) was cloned into a vector encoding an N-terminal $His_6$-SUMO tag between NdeI and XhoI restriction sites (subcloned by GenScript). Cells were re-suspended in lysis buffer (25 mM Tris, pH 8.0, 500 mM NaCl, 10 mM imidazole, and 2 mM TCEP) and sonicated for 15 min. The lysate was centrifuged at 10,000 rpm for 20 min at 4 °C to remove cellular debris. The supernatant was applied onto a HisTrap FF column and eluted using a linear gradient of 0–500 mM imidazole. Fractions with >90% purity were pooled and incubated with SUMO protease overnight at 4 °C to remove the $His_6$-SUMO tag, followed by dialysis into lysis buffer. The cleavage mixture was reapplied to a HisTrap FF column, and the flow-through containing the cleaved protein was further purified by size exclusion chromatography (HiLoad 16/600 Superdex 75 pg) in 25 mM Tris. pH 8.0, 150 mM NaCl, and 2 mM TCEP. Fractions of >95% purity estimated by SDS-PAGE, were pooled and buffer-exchanged into 25 mM HEPES, pH 7.4, 150 mM NaCl, and 2 mM TCEP and stored at –80° C until use.

**Isothermal titration calorimetry.** Titrations were performed on a Malvern MicroCal VP-ITC at 25.0 °C essentially as described earlier (3,4). Unless specified otherwise, 600 μM nucleotide was titrated into 50 μM *Ab*ZigA and the heat change was measured following 30 injections. The system was allowed 3 min to equilibrate between each injection and the stirring rate was set to 300 rpm. The heat change of the system was plotted against the molar ratio of nucleotide:protein and the data was fitted to a single binding site model using software provided by the instrument manufacturer. The first data point was omitted from all fittings. Calculated parameters for stoichiometry, $\Delta H$, and $K_a$ of the interaction were recorded with $n$ allowed to float, except for GTP into R293A/K295A and R293A *Ab*ZigA in which $n$ was fixed to 1.0 to allow fitting for $K_a$. ITC buffer consisted of 25 mM HEPES pH 7.4, 150 mM NaCl, 2 mM $MgCl_2$, and 2 mM TCEP. Ligand into buffer controls were performed to subtract heats of dilution from experimental datasets. Mutant *Ab*ZigA concentrations were adjusted to obtain better *c*-values when needed, with nucleotide concentrations adjusted to be 10-fold the initial protein concentration. 150 μM of K295A *Ab*ZigA was used for titrations with GTP while 200 μM R293A/K295A *Ab*ZigA and 150 μM R293A *Ab*ZigA used for titrations with GDP.

**Quin-2 Competition Assays.** These experiments were carried out essentially as described in our prior work (3,5,6), and the data globally fitted with a chelator competition model running in DynaFit (7).

**NMR Spectroscopy.** Uniformly $^{15}N$, $^{13}C$-labeled MigC core CTD (residues 233-332) was expressed in *E. coli* BL21 (DE3) cells in M9 minimal medium containing 1.0 g of $^{15}NH_3Cl$ and 2.0 g $^{13}C_6$-glucose as the sole nitrogen and carbon sources, respectively. Further expression, isolation and purification of these isotope-labeled proteins were performed as described above for unlabeled protein. NMR spectra were recorded at 30 °C on a 600 MHz or 800 MHz Bruker Avance Neo spectrometer equipped with a cryogenic probe in the METACyt Biomolecular NMR Laboratory at Indiana University, Bloomington. NMR samples for backbone resonance assignments contained 0.26 mM $^{15}N$,$^{13}C$-labeled MigC core CTD in 25 mM HEPES pH 7.0, 150 mM NaCl, 2 mM TCEP and 10% v/v $D_2O$, with 0.3 mM DSS as an internal reference. Backbone chemical shifts were assigned for each state using the following standard triple-resonance experiments: CBCACONH, HNCA, HNCACB, HNCACO, HNCO, HNCOCA and HBHACONH using non-uniform sampling with Poisson gap schedules. Data were acquired using Topspin 4.5.0 (Bruker) and processed using NMRPipe and istHMS and analyzed using CARA and NMRFAM-Sparky (8), running on NMRBox (9). TALOS-N was used for chemical shift-based secondary structure predictions and residue-specific $S^2$ order parameters (10).

Uniformly $^{15}N$-labeled MigC WT and RK mutant were expressed and purified as described above. Longitudinal ($T_1$) and transverse ($T_2$) relaxation times were measured for both the WT and RK MigC using TROSY-adapted standard HSQC relaxation sequences. $T_1$ and $T_2$ values were extracted using NMRFAM-Sparky (8) and then used to calculate the $T_1/T_2$ for each residue. These experimental values were compared to hydrodynamic predictions obtained from HYDRONMR (11) simulations based on the corresponding AF3 models. For inter-proton distances and tertiary $\beta$-sheet structure integrity, $^{15}N$-edited NOESY-HSQC spectra were recorded at 30 ºC on the 800 MHz instrument with a mixing time of 240 ms. NOESY spectra were processed using NMRpipe (12) and analyzed in Sparky (8) to identify inter-residue NOE contacts used for qualitative validation of the MigC core CTD structure.

NMR experiments to monitor binding of labeled MigC linker-core CTD (residues 207-332) to unlabeled full-length WT MigC or RK MigC were performed using uniformly $^{15}N$-labeled MigC linker-core CTD domains. The initial concentration of each was 0.35 mM in buffer described above. WT MigC or RK MigC was added into the solution of CTD to final concentration 0.6 mM. A $^{1}H$-$^{15}N$ HSQC spectrum was acquired at 30 °C on the 800 MHz spectrometer. The concentration of MigC CTD decreased from 0.35 mM to approximately 0.108 mM upon WT or RK addition. To maintain as approximately constant signal-to-noise ratio across spectra, the number of scans was scaled inversely with the concentration of MigC CTD. Residues showing substantial intensity loss were interpreted as undergoing intermediate conformational exchange as a result of complex formation.

**Generation of Bacterial Strains.** The strains and plasmids used in this study are listed in Table S4. The markerless *A. baumannii ΔmigC* mutant was generated by amplifying 1,500 bp DNA from the 5' and 3' flanking regions of *migC* from *A. baumannii* ATCC 17978VU gDNA and cloned into pFLP2 digested with XbaI and BamHI-HF using HiFi Assembly. Introduction of the construct into *A. baumannii* and subsequent allelic exchange was performed as previously described (2). The *migC* expression vectors were generated by amplifying the *migC* locus with its native promoter from *A. baumannii* ATCC 17978VU gDNA via PCR and cloned into pWH1266 digested with SalI and BamHI-HF using HiFi Assembly. Mutations were introduced into *migC* by amplifying *migC* and its native promoter from *A. baumannii* ATCC 17978VU gDNA using PCR primers that would result in two overlapping mutant fragments possessing either R274A, K276A or R274AK276A. The PCR fragments were then similarly cloned into SalI and BamHI-HF digested pWH1266 as described above. All resulting pWH1266 *migC* expression vectors were maintained in *E. coli* DH5α and transformed into the appropriate *A. baumannii* strain. *ΔzigA* complementation strains were similarly constructed by PCR amplifying the *zigA* locus with its native promoter from *A. baumannii* ATCC 17978VU gDNA using primers that would result in two overlapping mutant fragments possessing either R293A, K295A, or R293AK295A. The PCR fragments were cloned into pKNOCK-mTn7-Amp digested with KpnI and BamHI-HF using HiFi Assembly. The resulting wild-type or RxK pKNOCK-mTn7-Amp-*zigA* complementation constructs were transformed into *E. coli* DH5αλpir$^{+}$ and integrated into the chromosome of *A. baumannii ΔzigA* using *E. coli* HB101 (pRK2013) and *E. coli* EC100D (pTNS2) as previously described (13).

**Bacterial Growth Curves.** Overnight cultures from isolated colonies of each strain were back diluted 1:1000 into fresh LB and grown at 37°C shaking for 1 hr. These back dilutions were then used to inoculate 96-well microtiter plates at a 1:50 dilution into LB with or without 75 ug/ml carbenicillin (FisherScientific, BP2648250), or TPEN (*N,N,N′,N′*-tetrakis(2-pyridinylmethyl)-1,2-ethanediamine) (ThermoFischer, J64206-MC). Plates were grown at 37 °C with continuous shaking in a BioTek Epoch2 plate reader with the $OD_{600}$ recorded every 60 min.

### Supplementary Tables

**Table S1.** Thermodynamic parameters for the binding of G-nucleotides to RxKG mutant *Ab*ZigAs.[a]

| **Protein** | **Ligand** | $K_a$ ($M^{-1}$) | $\Delta H$ (kcal $mol^{-1}$) | $-T\Delta S$ (kcal $mol^{-1}$) | $\Delta G$ (kcal $mol^{-1}$) | $n$ |
|---|---|---|---|---|---|---|
| WT[b] | GTP | $2.8 \pm 0.5 \times 10^5$ | -24.5 ± 0.4 | 16.5 ± 0.3 | -8.0 ± 0.7 | 0.9 ± 0.10 |
| R293A/K295A | GTP | $4.4 \pm 0.1 \times 10^3$ | n.d.[c] | n.d.[c] | -5.0 ± 0.1 | 1[d] |
| R293A | GTP | $1.7 \pm 0.1 \times 10^4$ | n.d.[c] | n.d.[c] | -5.8 ± 0.2 | 1[d] |
| K295A | GTP | $1.3 \pm 0.3 \times 10^5$ | -30.3 ± 0.8 | 23.3 ± 0.9 | -7.0 ± 0.1 | 0.8 ± 0.04 |
| WT[b] | GDP | $8.8 \pm 0.6 \times 10^5$ | -27.3 ± 0.6 | 19.1 ± 0.7 | -8.1 ± 0.1 | 1.0 ± 0.04 |
| R293A/K295A | GDP | $5.5 \pm 0.2 \times 10^4$ | -20.4 ± 0.1 | 14.0 ± 0.1 | -6.5 ± 0.1 | 0.9 ± 0.03 |
| R293A | GDP | $1.3 \pm 0.3 \times 10^5$ | -20.4 ± 0.9 | 13.8 ± 0.8 | -6.6 ± 0.1 | 1.0 ± 0.02 |
| K295A | GDP | $6.0 \pm 0.1 \times 10^5$ | -28.9 ± 0.1 | 21.1 ± 0.1 | -7.9 ± 0.1 | 0.9 ± 0.07 |

[a]From duplicate experiments. $n$, stoichiometry; $K_a$, association equilibrium constant. Conditions: 25 mM HEPES pH 7.4, 150 mM NaCl, 2 mM $MgCl_2$, 2 mM TCEP, 25.0 °C. [b]Taken from Osterberg *et al.* (3). [c]Not determined due to a low $K_a$. [d]Stoichiometry fixed during ITC analysis.

**Table S2.** GTP hydrolysis rates for RxKG mutant *Ab*ZigA and *Ab*MigC enzymes measured in this study[a]

| **Enzyme** | **Metal status** | **CTD[b]** | **Rep 1, $V_o$[c]** | **Rep 2, $V_o$[c]** | **Mean $V_o$** | **STDEV mean $V_o$** |
|---|---|---|---|---|---|---|
| ZigA WT | Apo | - | 0.0316 | 0.0252 | 0.0284 | 0.0045 |
| ZigA WT | Zn | - | 0.106 | 0.105 | 0.106 | 0.001 |
| ZigA R293A/K295A | Apo | - | 0.0036 | 0.0065 | 0.0051 | 0.0021 |
| ZigA R293A/K295A | Zn | - | 0.0054 | 0.0041 | 0.0047 | 0.0009 |
| ZigA R293A | Apo | - | 0.0026 | 0.0123 | 0.0074 | 0.0066 |
| ZigA R293A | Zn | - | 0.0070 | 0.0056 | 0.0063 | 0.0010 |
| ZigA K295A | Apo | - | 0.0151 | 0[d] | 0.0075 | 0.0107 |
| ZigA K295A | Zn | - | 0.0048 | 0[d] | 0.0024 | 0.0056 |
| MigC WT | Apo | - | 0.203 | 0.137 | 0.170 | 0.047 |
| MigC WT | Zn | - | 0.676 | 0.718 | 0.697 | 0.029 |
| MigC R274A/K276A | Apo | - | 0.0051 | 0.0046 | 0.0049 | 0.0004 |
| MigC R274A/K276A | Zn | - | 0.0052 | 0.0013 | 0.0033 | 0.0027 |
| MigC R274A | Apo | - | 0.0058 | 0.0099 | 0.0079 | 0.0030 |
| MigC R274A | Zn | - | 0.0118 | 0.0143 | 0.0131 | 0.0018 |
| MigC K276A | Apo | - | 0.0099 | 0.0159 | 0.0130 | 0.0042 |
| MigC K276A | Zn | - | 0.0454 | 0.0486 | 0.0470 | 0.0022 |
| ZigA R293A/K295A | Apo | ZigA CTD | 0.0066 | 0.0086 | 0.0076 | 0.0014 |
| ZigA R293A/K295A | Zn | ZigA CTD | 0.0634 | 0.0877 | 0.0756 | 0.0172 |
| ZigA R293A/K295A | Apo | MigC CTD | 0.0070 | 0[d] | 0.0031 | 0.0056 |
| ZigA R293A/K295A | Zn | MigC CTD | 0.0066 | 0.0022 | 0.0044 | 0.0031 |
| MigC R274A/K276A | Apo | MigC CTD | 0.0804 | 0.0806 | 0.0805 | 0.0001 |
| MigC R274A/K276A | Zn | MigC CTD | 0.801 | 0.764 | 0.782 | 0.026 |
| MigC R274A/K276A | Apo | ZigA CTD | 0.0026 | 0.0316 | 0.0171 | 0.0205 |
| MigC R274A/K276A | Zn | ZigA CTD | 0.0074 | 0.0054 | 0.0064 | 0.0014 |

| | | | | | | |
|---|---|---|---|---|---|---|
| ZigA E258A | Apo | - | 0.0725 | 0.0781 | 0.0754 | 0.0039 |
| ZigA E258A | Zn | - | 0.0696 | 0.0622 | 0.0659 | 0.0052 |
| ZigA R293A/K295A | Apo | ZigA core CTD | 0.263 | 0.195 | 0.228 | 0.048 |
| ZigA R293A/K295A | Zn | ZigA core CTD | 0.241 | 0.212 | 0.227 | 0.021 |
| ZigA R293A/K295A | Apo | *Pa*ZigA CTD | 0.156 | 0.144 | 0.150 | 0.008 |
| ZigA R293A/K295A | Zn | *Pa*ZigA CTD | 0.151 | 0.144 | 0.147 | 0.005 |
| MigC R274A/K276A | Apo | MigC core CTD | 0.0225 | 0.0617 | 0.0421 | 0.0277 |
| MigC R274A/K276A | Zn | MigC core CTD | 1.132 | 0.968 | 1.050 | 0.115 |

[a]Using the malachite green-based kinetic assay with other details described in Materials and Methods. Data presented in graphical form in Fig. 4, main text. [b]Unless otherwise indicated CTD corresponds to linker-core CTD present at 25 μM. [c]Units are μmol $P_i$ min$^{-1}$ μmol enzyme$^{-1}$. [d]The lower limit for this assay is ≤0.0002 μmol $P_i$ min$^{-1}$ μmol enzyme$^{-1}$.

**Table S3.** Zinc binding affinities ($K_{Zn}$) of mutant ZigA enzymes[a]

| *Ab*ZigA variant | $K_{Zn}$ ($M^{-1}$) | $K_{Zn}$, +GTPγS ($M^{-1}$) | $\Delta G_c$ (kcal $mol^{-1}$)[d] |
|---|---|---|---|
| WT[b] | 8.7 ±0.5 x $10^{10}$ | 3.2 ±0.4 x $10^{12}$ | –2.1 ±0.2 |
| RK | 8.2 ±0.8 x $10^{10}$ | 4.4 ±0.5 x $10^{11}$ | –1.0 ±0.1 |
| E258 | 4.9 ±0.8 x $10^{11}$ | n.d.[c] | n.d. |

[a]Determined using quin-2 competition assays like those shown in Figure S3. The mean and standard deviation obtained from a global fit of two independent experiments are shown. [b]From ref (3). [c]n.d., not determined. [d]From $\Delta G_c = -RT\ln(K_{Zn\ +GTP\gamma S}/K_{Zn})$. Conditions: 25 mM Hepes, 150 mM NaCl, 2 mM $MgCl_2$, 25.0 ºC.

**Table S4**. Bacterial strains, plasmids and oligonucleotide primers used in this study

| **Strain** | **Source** | **Description** |
|---|---|---|
| *Acinetobacter baumannii* ATCC 17978VU | ATCC (14) | Wild-type |
| *A. baumannii* ATCC 17978VU Δ*migC* | This work | Contains deletion of the *migC* gene |
| *A. baumannii* ATCC 17978VU Δ*migC*:*migC* | This work | Δ*migC* complemented with wild-type *migC* under control of its own promoter |
| *A. baumannii* ATCC 17978VU Δ*migC*:*migC*$^{R274A/K276A}$ | This work | Δ*migC*complemented with a R264A/K276A mutant *migC* allele under control of its own promoter |
| *A. baumannii* ATCC 17978VU Δ*migC*:*migC*$^{K276A}$ | This work | Δ*migC* complemented with a K276A mutant *migC* allele under control of its own promoter |
| *A. baumannii* ATCC 17978VU Δ*migC*:*migC*$^{R274A}$ | This work | Δ*migC* complemented with a R274A mutant *migC* allele under control of its own promoter |
| *A. baumannii* ATCC 17978VU Δ*zigA* | (15) | Contains deletion of the *zigA* gene |
| *A. baumannii* ATCC 17978VU Δ*zigA*:*zigA* | This work | Δ*zigA* chromosomally complemented with wild-type *zigA* under control of its own promoter |
| *A. baumannii* ATCC 17978VU Δ*zigA*:*zigA*$^{R293A/K295A}$ | This work | Δ*zigA* chromosomally complemented with a R293A/K295A mutant *zigA* allele under control of its own promoter |
| *A. baumannii* ATCC 17978VU Δ*zigA*:*zigA*$^{K295A}$ | This work | Δ*zigA* chromosomally complemented with a K295A mutant *zigA* allele under control of its own promoter |
| *A. baumannii* ATCC 17978VU Δ*zigA*:*zigA*$^{R293A}$ | This work | Δ*zigA* chromosomally complemented with a R293A mutant *zigA* allele under control of its own promoter |
| *E. coli* DH5α pWH1266 | This work | Empty complementation vector control |
| *E. coli* DH5α pWH1266 -*migC* | This work | Wild-type *migC* complementation vector |
| *E. coli* DH5α pWH1266 -*migC*$^{R274A}$ | This work | *migC*$^{R274A}$ complementation vector |
| *E. coli* DH5α pWH1266-*migC*$^{K276A}$ | This work | *migC*$^{K276A}$ complementation vector |
| *E. coli* DH5α pWH1266-*migC*$^{R274A/K276A}$ | This work | *migC*$^{R274A/K276A}$ complementation vector |

| | | |
|---|---|---|
| *E. coli* DH5αλpir⁺ pKNOCK-mTn7-*zigA* | This work | Wild-type *zigA complementation vector* |
| *E. coli* DH5αλpir⁺ pKNOCK-mTn7-*zigA*$^{R293A}$ | This work | *zigA*$^{R293A}$ complementation vector |
| *E. coli* DH5αλpir⁺ pKNOCK-mTn7-*zigA*$^{K295A}$ | This work | *zigA*$^{K295A}$ complementation vector |
| *E. coli* DH5αλpir⁺ pKNOCK-mTn7-*zigA*$^{R293A/K295A}$ | This work | *zigA*$^{R293A/K295A}$ complementation vector |
| *E. coli HB101 pRK2013* | (16) | *E. coli* carrying the mobilization helper plasmid |
| *E. coli* pTNS2 | (17) | Encodes transposase required for *A. baumannii* chromosomal complementation |
| pFLP2 | (18) | Allelic exchange vector for *A. baumannii* |
| pWH1266 | (19) | Complementation expression plasmid for *A. baumannii* |
| pWH1266-*migC* | This work | Expression plasmid encoding for *migC* regulated by the native promoter of *migC* |
| pWH1266-*migC*$^{R274A}$ | This work | Expression plasmid encoding for a R274A mutant allele of *migC* regulated by the native promoter of *migC* |
| pWH1266-*migC*$^{K276A}$ | This work | Expression plasmid encoding for a K276A mutant allele of *migC* regulated by the native promoter of *migC* |
| pWH1266-*migC*$^{R274A/K276A}$ | This work | Expression plasmid encoding for a R274A/K276A mutant allele of *migC* regulated by the native promoter of *migC* |
| pKNOCK-mTn7-*Amp*-*zigA*$^{R293A/K295A}$ | This work | Complementation vector containing a R293A/K295A mutant allele of *zigA* regulated by the native promoter of *zigA* |
| pKNOCK-mTn7-*Amp*-*zigA*$^{R293A}$ | This work | Complementation vector containing a R293A mutant allele of *zigA* regulated by the native promoter of *zigA* |
| pKNOCK-mTn7-Amp-*zigA*$^{K295A}$ | This work | Complementation vector containing a K295A mutant allele of *zigA* regulated by the native promoter of *zigA* |
| **Primer** | **Sequence** | **Description** |
| AD01_pFLP2_migc_UP_F | ggttaaaaaggatcgatcctttcgtcatacca atagctgttatttc | 5' flanking region for *migC* KO construct using pFLP2, forward |

| | | |
|---|---|---|
| AD02_pFLP2_migc_UP_R | tcaatgttctaagcggctcaaatcaacaag | 5’ flanking region for *migC* KO construct using pFLP2, reverse |
| AD03_pFLP2_migc_DOWN_F | tgagccgcttagaacattgaaatacttcaatattctctatgagag | 3’ flanking region for *migC* KO construct using pFLP2, forward |
| AD04_pFLP2_migc_DOWN_R | aagttcctattctctaggggccgggcgtcggtatgaatatc | 3’ flanking region for *migC* KO construct using pFLP2, reverse |
| AD05_migc_external_UP | gaacctcatataaaacgatgctgtgtc | Upstream *migC* screening primer |
| AD06_migc_external_DOWN | ctttaagcgccatagttaatatgtcca | Downstream *migC* screening primer |
| AD07_pFLP2_F | tgaacggcaggtatatgtgatggg | pFLP2 sequencing primers, forward |
| AD08_pFLP2_R | aagcgctcgttttcggaaacg | pFLP2 sequencing primers, reverse |
| AD11_pKNOCK_zigA_R293A_F | gggagtcgtggcgtccaaaggctttttc | To generate *zigA*$^{R293A}$ under native *zigA* promoter for pKNOCK complementation construct, forward |
| AD12_pKNOCK_zigA_R293A_R | gaaaaagcctttggacgccacgactccc | To generate *zigA*$^{R293A}$ under native *zigA* promoter for pKNOCK complementation construct, reverse |
| AD13_pKNOCK_zigA_K295A_F | cgtgcgttccgcgggctttttctg | To generate *zigA*$^{K295A}$ under native *zigA* promoter for pKNOCK complementation construct, forward |
| AD14_pKNOCK_zigA_K295A_R | cagaaaaagcccgcggaacgcacg | To generate *zigA*$^{K295A}$ under native *zigA* promoter for pKNOCK complementation construct, reverse |
| AD15_pKNOCK_zigA_R293A_K295A_R | gccaaccagaaaaagcctgcggaagccacgactcccggccattc | To generate *zigA*$^{R293A/K295A}$ under native *zigA* promoter for pKNOCK complementation construct, reverse |
| AD16_pKNOCK_zigA_R293A_K295A_F | gaatggccgggagtcgtggcttccgcaggctttttctggttggc | To generate *zigA*$^{R293A/K295A}$ under native *zigA* promoter for pKNOCK complementation construct, forward |
| AD39_zigA_pKNOCK F | catgcatgagctcactagtgaacactaagtgatttggc | Clone wild-type *zigA* under native promoter for pKNOCK complementation construct, forward |
| AD40_zigA_pKNOCK R | gcaaggccttcgcgaggtacttaagcgatcaacatcgtttc | Clone wild-type *zigA* under native promoter for pKNOCK complementation construct, reverse |
| AD43_pKNOCK_seq F | gcgcttttgaagctaattcg | pKNOCK insert screening primer, forward |
| AD44_pKNOCK_seq R | atttcacttatctggttggcc | pKNOCK insert screening primer, reverse |
| AD45_pKNOCK_Tn7_screen F | tatggaagaagttcaggctcgtggcgg | Tn7 insertion screening primer, forward |
| AD46_pKNOCK_Tn7_screen R | cacagcataactggactgatttc | Tn7 insertion screening primer, reverse |

| | | |
|---|---|---|
| AD37_ pWH1266_migC_F | gcgaccacacccgtcctgtgcaagttagttacgtaaacaaaac | Clone wild-type *migC* under native promoter for pWH1266 expression construct, forward |
| AD38_ pWH1266_migC_R | aaggctctcaagggcatcggctattgttcaattctgcaag | Clone wild-type *migC* under native promoter for pWH1266 expression construct, reverse |
| AD21_ pWH1266_migC_R274A_F | agactggctagcgattaagggaatttttaatac | To generate *migC*$^{R274A}$ under native *migC* promoter for pWH1266 expression construct, forward |
| AD22_ pWH1266_migC_R274A_R | gtattaaaaattcccttaatcgctagccagtct | To generate *migC*$^{R274A}$ under native *migC* promoter for pWH1266 expression construct, reverse |
| AD23_ pWH1266_migC_K276A_F | gctacgtattgcgggaatttttaatacag | To generate *migC*$^{K276A}$ under native *migC* promoter for pWH1266 expression construct, forward |
| AD24_ pWH1266_migC_K276A_R | ctgtattaaaaattcccgcaatacgtagc | To generate *migC*$^{K276A}$ under native *migC* promoter for pWH1266 expression construct, reverse |
| AD25_ pWH1266_migC_R274A_K276A_R | gatctgtattaaaaattcccgcaatagctagccagtcttgttgtgcacacaacacatcgag | To generate *migC*$^{R274A/K276A}$ under native *migC* promoter for pWH1266 expression construct, reverse |
| AD26_ pWH1266_migC_R274A_K276A_F | ctcgatgtgttgtgtgcacaacaagactggctagctattgcgggaatttttaatacagatc | To generate *migC*$^{R274A/K276A}$ under native *migC* promoter for pWH1266 expression construct, forward |
| *MigC_pHIS_R* | tcgacgtaggcctttgaattcctattgttcaattctgcaagctaata | Cloning *a1s_0934* into pHIS-Parallel1, reverse |
| *MigC_R274A_pHIS_F* | cacaacaagactggctagcgattaagggaatttttaatacagatcagg | To generate R274A point mutant construct in pHIS- Parallel1, forward |
| *MigC_K276A_pHIS_F* | cacaacaagactggctacgtattgcgggaatttttaatacagatcagg | To generate K276A point mutant construct in pSUMO, forward |
| *MigC_R274AK276A_pHIS_F* | cacaacaagactggctagcgattgcgggaatttttaatacagatcagg | To generate R274A/K276A double mutant construct in pHIS- Parallel1, forward |
| *MigC_R274A-K276A_pHIS_R* | cacacaacacatcgagcaaagc | To generate K276A, R274A, and R274A/K276A point mutants construct in pSUMO, reverse |
| *MigC_CTD_F* | cgaacagattggaggatgcactgaacggtctatacaacctttac | Cloning MigC CTD into pSUMO, forward |
| *ZigA_pHIS_R293AK295A_F* | tttaactttaagaaggagatatacatatgatgcacactgcaatcgcca | To generate R293A/K295A double mutant construct in pHIS- Parallel1, forward |

| | | |
|---|---|---|
| *ZigA_pHIS_R293AK295A_R* | ggtgatggtagtacgacattaagcgatcaacatcgtttcatcatcttg | To generate R293A/K295A double mutant construct in pHIS- Parallel1, reverse |
| *ZigA_pHIS_R293A_F* | gggagtcgtggcgtccaaaggctttttc | To generate R293A point mutant construct in pHIS- Parallel1, forward |
| *ZigA_pHIS_R293A_R* | ggccattcagcctgtac | To generate R293A point mutant construct in pHIS- Parallel1, reverse |
| *ZigA_pHIS_K295A_F* | cgtgcgttccgcgggctttttctg | To generate K295A point mutant construct in pHIS- Parallel1, forward |
| *ZigA_pHIS_K295A_R* | actcccggccattca | To generate K295A point mutant construct in pHIS- Parallel1, reverse |
| *ZigA_pHIS_E258A_F* | gcatacacctgcgactgaagagtatg | To generate E258A point mutant construct in pHIS- Parallel1, forward |
| *ZigA_pHIS_E258A_R* | tcgccgcgtaactct | To generate E258A point mutant construct in pHIS- Parallel1, reverse |
| *PaZigA_pHis_F* | gtttaactttaagaaggagatatacatATGAACCAACGCCTACCTGTC | Cloning PAO1_*PA5535* into pHIS-Parallel1, forward |
| *PaZigA_pHis_R* | ggtgatggtagtacgacatTCATGCGGCGTGCCCCCAGTC | Cloning PAO1_*PA5535* into pHIS-Parallel1, reverse |

### Supplementary Figures

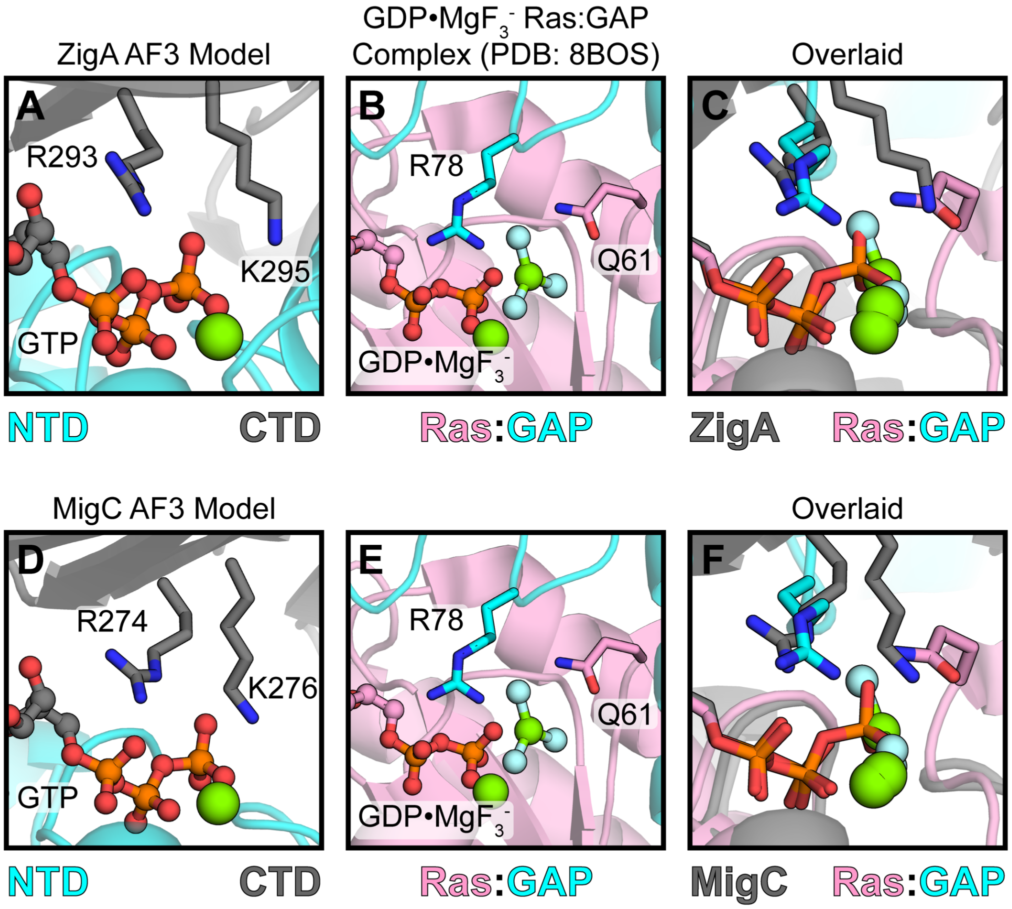


**Fig. S1.** Expanded regions of the structural models of one selected Ras-GAP complex bound to the GTP analogs compared to AF3-derived ZigA and MigC "closed" conformations modeled in the presence of Mg(II)•GTP. (*A*) Expanded view of of *Ab*ZigA showing the CTD (grey) RxKG motif in proximity to the NTD (cyan) nucleotide binding pocket. (*B*) Expanded view of Ras (pink) bound to transition state analog Mg(II)•GDP-$AlF_3$ in complex with its GAP effector protein (cyan) (PDB: 8BOS). (*C*) Overlay of structures shown in panels A and B. (*D*) Expanded view of of *Ab*MigC showing the CTD (grey) RxKG motif in proximity to the NTD (cyan) nucleotide binding pocket. **(***E***)** Same as panel B. (*F*) Overlaid of the structures shown in panels D and E.

| | cluster 1 | | | | cluster 3 | | cluster 4 | cluster 7 | cluster 26 | |
|---|---|---|---|---|---|---|---|---|---|---|
| | *Ab*ZigA | *Rj*Nha3 | *Bs*ZagA | *Pa*ZigA | *Ec*YjiA | *Hs*ZNG1 | *Rc*CobW | *Ec*YeiR | *Ab*MigC | |
| *Ab*ZigA | | 68.4 | 69.2 | 74.5 | 50.5 | 44.1 | 54.0 | 41.9 | 42.5 | histidine catabolism; cell wall homeostasis |
| *Rj*Nha3 | 54.2 | | 67.2 | 73.8 | 44.8 | 43.5 | 51.7 | 43.5 | 44.9 | Fe-nitrile hydratase maturation |
| *Bs*ZagA | 57.8 | 51.3 | | 65.8 | 46.0 | 43.9 | 48.6 | 38.3 | 41.0 | folate biosynthesis |
| *Pa*ZigA | 60.8 | 61.3 | 49.9 | | 46.7 | 44.3 | 54.6 | 44.7 | 46.7 | unknown |
| *Ec*YjiA | 28.6 | 24.4 | 26.0 | 24.1 | | 48.3 | 51.7 | 38.4 | 39.0 | unknown |
| *Hs*ZNG1 | 23.5 | 21.8 | 21.9 | 23.5 | 27.6 | | 48.0 | 43.8 | 40.1 | methionine aminopeptidase-I metalation |
| *Rc*CobW | 27.0 | 28.2 | 25.0 | 27.3 | 31.1 | 27.0 | | 45.9 | 43.1 | cobalamin biogenesis |
| *Ec*YeiR | 21.6 | 20.7 | 18.2 | 21.6 | 21.6 | 21.9 | 23.7 | | 55.0 | unknown |
| *Ab*MigC | 19.9 | 21.4 | 17.5 | 24.1 | 18.7 | 21.1 | 22.3 | 35.3 | | cell wall biogenesis or homeostasis |

**Fig. S2.** Pairwise sequence identity (below diagonal) and similarity (above diagonal) matrix of all the COG0523 enzymes discussed in this work (see Fig. 1*C-E*, main text). The SSN cluster designation from Edmonds *et al.* (20) is shown above the matrix, with insights into biological functions indicated. References to these functions are given in the text.

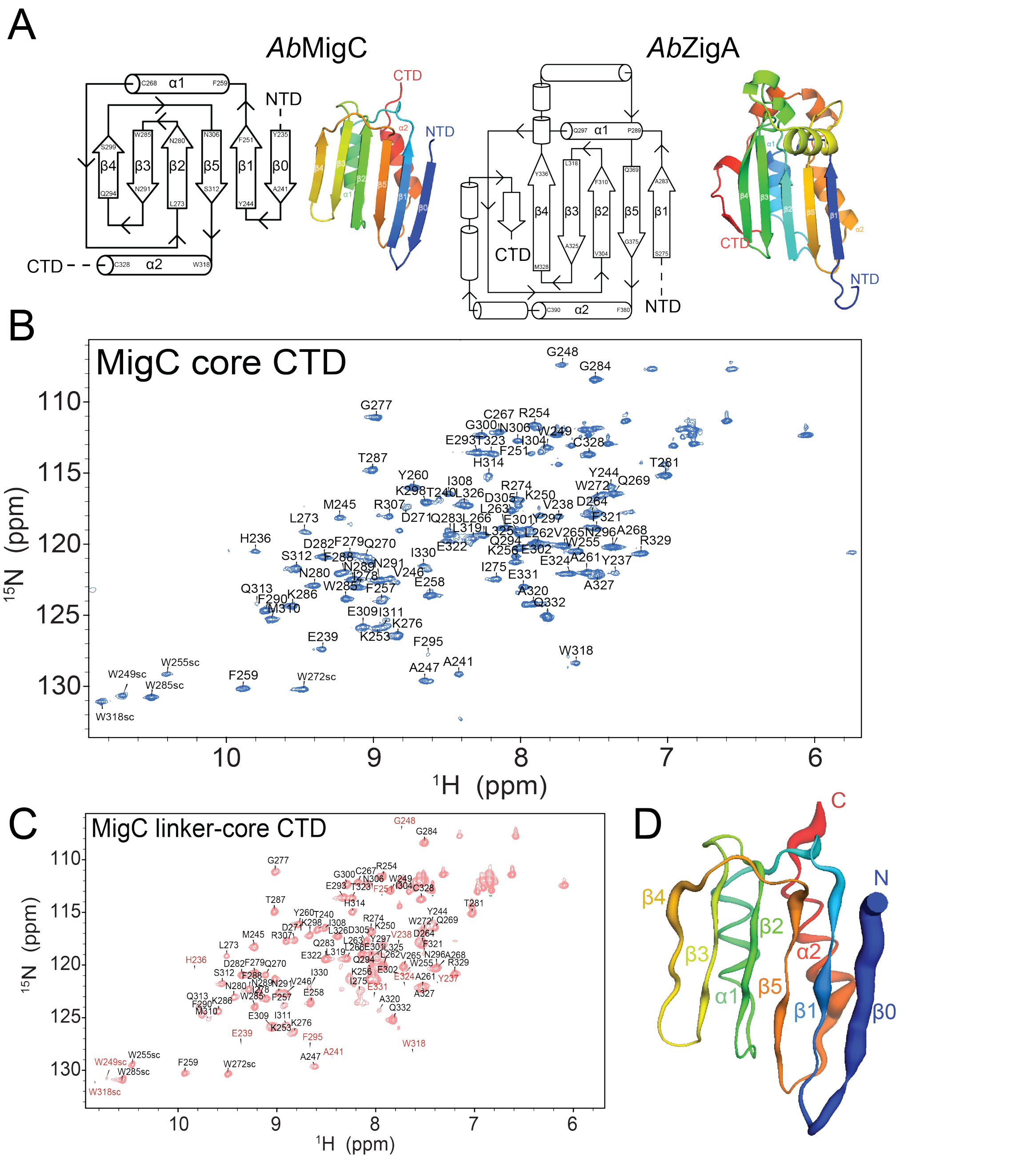


**Fig. S3.** (*A*) Secondary structural schematics of the tertiary structure (*left*) or AlphaFold3 models (*right*) of the C-terminal domains (CTDs) of *Ab*ZigA (ModelArchive deposition, ma-tca8p) and *Ab*MigC (ModelArchive, ma-iwqsx). The residue numbers defining secondary structure elements are indicated in each schematic. NTD, N-terminal G-domain (upstream). In the AF3 models (*right*), ribbons are shaded from *blue* to *red*, beginning with β1 (ZigA) or β0 (MigC), then proceeding through the C-terminal α2 helix (back). (*B*) Backbone $^{1}$H, $^{15}$N assignments of *Ab*MigC core CTD shown on the $^{1}$H,$^{15}$N HSQC spectrum of $^{13}$C,$^{15}$N-labeled *Ab*MigC core CTD (L233-Q332). (*C*) Backbone $^{1}$H,$^{15}$N assignments of *Ab*MigC core CTD shown on the $^{1}$H,$^{15}$N HSQC spectrum of $^{13}$C, $^{15}$N-labeled *Ab*MigC linker-core CTD (T207-Q332). Backbone crosspeaks broadened or missing from the linker-core CTD spectrum, but present in the core spectrum are highlighted in *red*. No attempt was made to assign the linker NH resonances, given their variable crosspeak intensities and poor chemical shift dispersion. **(*D*)** AF3 model of *Ab*MigC illustrating residues showing line broadening in the linker-core CTD $^{1}H,^{15}N$ HSQC spectrum vs. core CTD $^{1}H,^{15}N$ HSQC spectrum. The putty scale represents residue-specific line broadening of the amide resonance, with thicker segments indicating greater line broadening and thinner segments indicating minimal changes in peak intensity between the core CTD and linker-core CTD spectra. The ribbon is colored from *blue* (N) to *red* (C), beginning with $\beta$0, and proceeding through C-terminal $\alpha$2 helix (back). These data reveal that appending the N-terminal linker region to the MigC core domain gives rise to dynamical disorder localized to the $\beta$0, $\beta$1 and the N-terminal region of the $\alpha$2 helix in the AF3 model, all physically proximate to the appended linker.

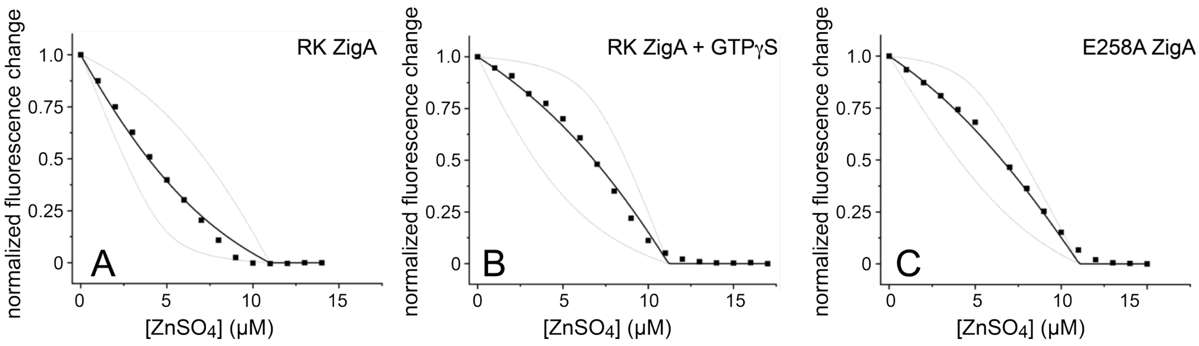


**Fig. S4.** Representative quin-2-metallochaperone Zn(II) binding competition curves for RK ZigA (*A*), RK ZigA + GTPγS (*B*) and E258A ZigA (*C*). The continuous line a fit a 1:1 competition model, with, $K_{Zn}$ for the enzyme optimized, with the dashed lines indicative of a $K_{Zn}$ value 10-fold higher or 10-fold lower than the fitted $K_{Zn}$ to illustrate the robustness of the fit. Globally fitted parameters from duplicate experiments are compiled in Table S3.

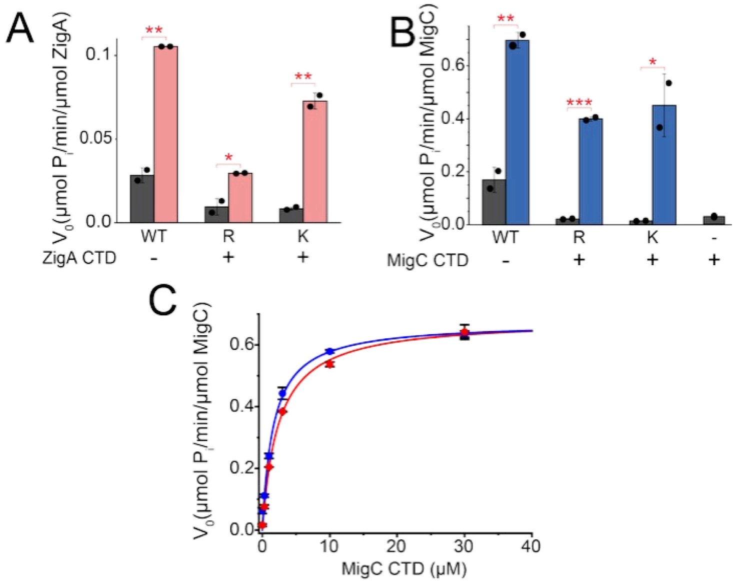


**Fig. S5.** Rescue of GTP hydrolysis activity of R and K mutants of ZigA. (*A*) and MigC (*B*) by cognate linker-core CTD. Linker-core CTD was present at 25 μM in all reactions with 5 μM enzyme in all cases, except for the last bar which was CTD alone with no enzyme (–). (*C*) MigC linker-core CTD-concentration dependence of the rescue of the R (*red* symbols) and the K (*blue* symbols) MigC mutants. The continuous lines through the data are fits to a hyperbolic model. Kinetic parameters for R MigC are $K_m$ = 2.4 ± 0.1 μM with $V_{max}$ = 0.68 ± 0.02 μmol $P_i$ $min^{-1}$ $\mu mol^{-1}$ MigC (*n*=2). Kinetic parameters for K MigC were determined $K_m$ = 1.66 ± 0.05 μM with $V_{max}$ = 0.674 ± 0.002 μmol $P_i$ $min^{-1}$ $\mu mol^{-1}$ MigC (*n*=2). •, $p<0.05$; **, $p<0.01$; ***, $p<0.001$.

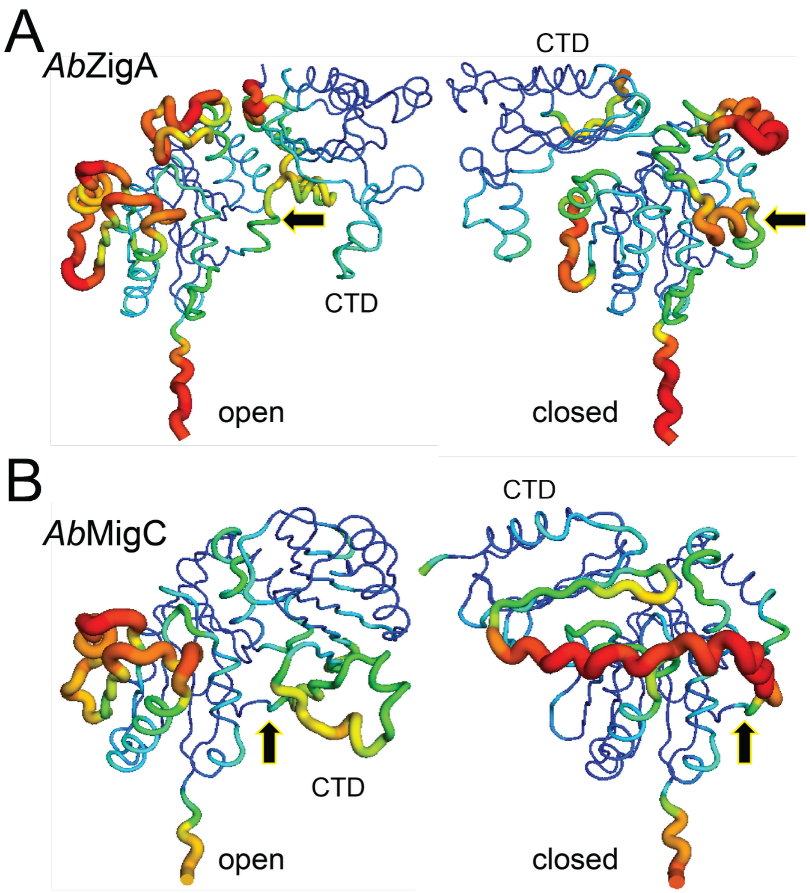


**Fig. S6.** Putty plots of the open and closed-state AF3 models for *Ab*ZigA (*A*) and *Ab*MigC (*B*) used in this study. In all cases, the chain is shaded from *blue* (high pLDDT) to *red* (low pLDDT) with the cartoon width changing from narrow (*blue*, well-modeled) to thick (*red*, poorly modeled). The N-terminus is pointing down, with the G-domains in a fixed position in all four models. No ligands are shown here, for clarity. *Black* arrow, the pivot point for CTD migration in each case. (*A*) Open, *Ab*ZigA bound to Zn(II) (ma-d36dv); closed, *Ab*ZigA bound to Mg(II)•GTP and Zn(II) (ma-tca8p). (*B*) Open, *Ab*MigC bound to Zn(II) (ma-l1a07); closed, *Ab*MigC bound to Mg(II)•GTP and Zn(II) (ma-iwqsx).

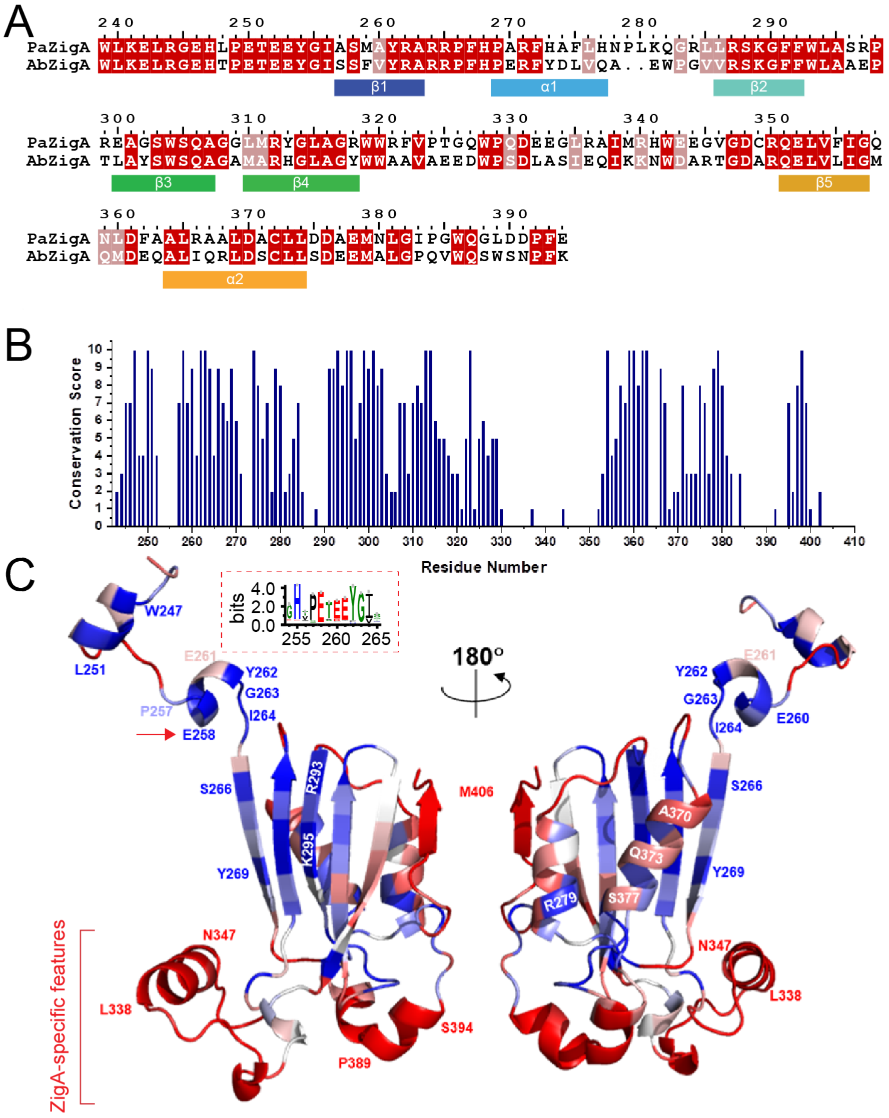


**Fig. S7**. Comparison of *Pseudomonas aeruginosa* (*Pa*) PAO1 ZigA (locus tag PA5535; uniprot ID Q9HT39, 400 residues) with *Ab*ZigA C-terminal domains. (*A*) Sequence alignment of the C-terminal domain of PaZigA and AbZigA, with the seven core secondary structural units (N-β1-α1-β2-β3-β4-β5-α2-C) highlighted below the sequences. The numbers above the sequence reflect the *Pa*ZigA residue numbers. (*B*) Sequence conservation score as a function of residue number position in AbZigA sequence (W247 is W239 in *Pa*ZigA) using 2266 sequences obtained from further fragmentation of the SSN cluster 1 sequence from ref. (20) using an alignment score of 140, and taking cluster 1 sequences from this analysis done by Jalview (Fig. S8). (*C*) Sequence conservation score painted on the AF3 model of *Ab*ZigA "closed" Mg•GTP bound state (ma-tca8p) with selected residues highlighted using residue numbers for *Ab*ZigA. ZigA-specific insertions, positioned at the bottom of these models are poorly conserved and thus may represent member-specific functionally important specificity determinants for apoenzyme client engagement and regulated metal transfer.

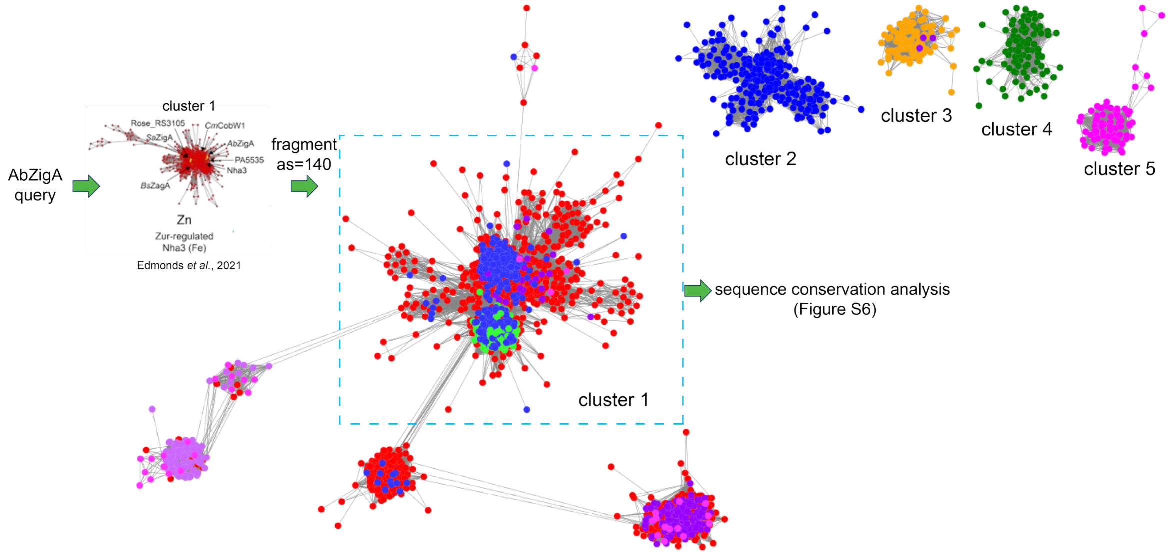


**Fig. S8**. Fragmentation of the SSN cluster 1 (20) sequences using an alignment score of 140, with the top five SSN clusters (by sequence count) shown for clarity. Sequences in the *cyan* box most closely related to *Ab*ZigA and *Pa*ZigA were used to generate the conservation score plot and model of *Ab*ZigA-like sequences (Fig. S7B-C). *Blue* nodes in main *red* (cluster 1) sequences correspond to those *zigA*-like genes that are genomically localized next to a PA5534 (DUF1826) family protein.

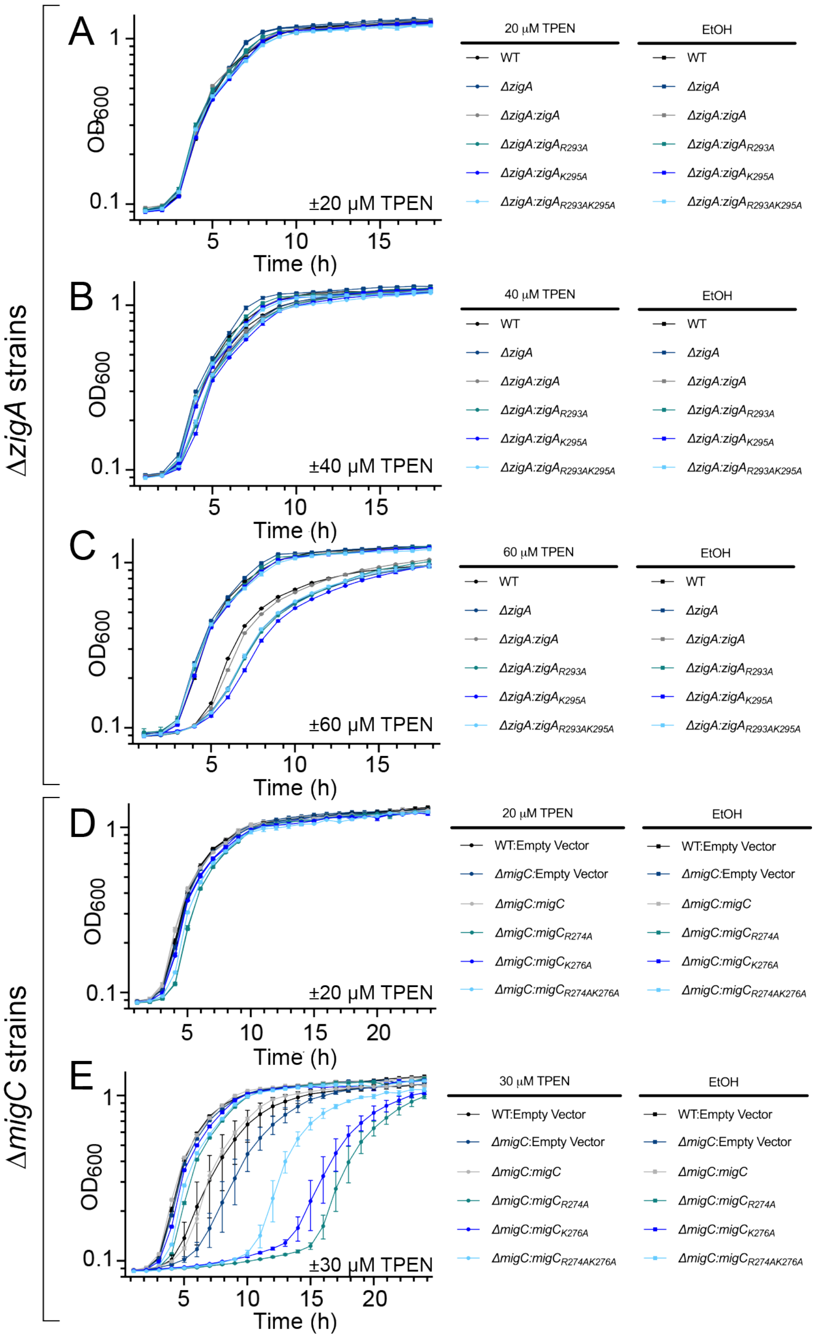


**Fig. S9**. Growth curves shown for the indicated *A. baumannii* strains under the conditions shown. (*A*)-(*C*) Δ*zigA* strains (see legend) chromosomally complemented with the indicated *zigA* allele compared to the parent wild-type (WT) strain grown in the presence of (*A*) 20 µM, (*B*) 40 µM or (*C*) 60 µM TPEN vs. the same amount of ethanol used to dissolve the TPEN (–TPEN). The same experiment conducted at 80 µM TPEN is shown in the main text (Fig. 6*A*). (*D*)-(*E*) Δ*migC* strains (see legend) harboring a pWH1266 complementation vector with the indicated *migC* allele compared to the parent wild-type (WT) strain grown in the presence of (*D*) 20 µM or (*E*) 30 µM TPEN vs. no addition of TPEN (–TPEN). The same experiment conducted at 40 µM TPEN is shown in the main text (Fig. 6*B*).
